# Forest biogeochemical monitoring indicates altered microbial communities, macronutrient availability, CO_2_ emissions and litter chemistry in root zone soil of oak trees with Acute Decline symptoms

**DOI:** 10.1101/2025.04.22.650028

**Authors:** Selvakumar Dhandapani, Thom McNeil, Bingyuan Lu, Oliver Booth, Luci Corbett, James Lunn, Adrian Nightingale, Xize Niu, Tiina Roose, Jiji John, Hanqing Lin, Adetunji Alex Adekanmbi, Liz J. Shaw

**Author notes:** **Corresponding Author**, Dr Selvakumar Dhandapani, Environment and Marine Sciences Division, Agri-Food Bioscience Institute (afbi), 18a Newforge lane, Belfast BT9 5PX, UK., Contact.

## Abstract

Acute and Chronic Oak Decline (AOD & COD) widely impact oak woodlands across Europe. AOD is known to cause rapid decline in tree health in a short span of 3-5 years. However, the interactions between such oak decline and soil biogeochemical dynamics are not fully understood. We selected three oak trees for each of the three treatments for biogeochemical monitoring: 1. No symptoms (Healthy), 2. AOD & 3. COD symptoms, in Writtle woodlands, Essex, UK. The selected trees were used for quarterly root zone soil sampling (cores to depth of 40 cm), and monthly surface soil (0-10 cm), leaf litter and GHG emissions sampling over a year (2022-23). Soil samples were characterised for their physico-chemical properties, total nutrient content & availability. Leaf litter samples were sorted into branches, leaves, seeds and flowers, weighed and their chemistry characterised. Soil microbial communities were characterised using phospholipid fatty acid analyses. Almost all the measured properties showed significant changes with season and depth, indicating a strong temporal and depth effect on forest soil biogeochemical dynamics. Notably macro-nutrient contents of leaf litter were greater in the summer period, which resulted in increased soil nutrient availability in autumn. This pattern was particularly exaggerated in AOD trees, which had greater nutrient content in litterfall in summer and greater nutrient availability in root zone soil in autumn than those of COD and Healthy trees. We found that AOD root zone soil have altered microbial communities, decreased CO_2_ emissions and increased macronutrient availability. Taken together, our study shows poor physiological nutrient management by diseased AOD trees revealed by greater nutrient concentration in AOD leaf litter indicating the lack of nutrient resorption and re-allocation before leaf senescence, and less active root zone soil under AOD trees, shown by reduced soil CO_2_ emissions and increased nutrient availability in oak root influenced zones. Hence, we conclude that the biogeochemical dynamics of AOD root zone soils are significantly different to that of healthy and COD trees, however controlled experiments are needed alongside these field observations to disentangle the causes from effects in understanding the relationship between soil health and oak decline.

**Highlights:**

- We monitored forest soil biogeochemical dynamics under oak trees of different health status
- Acute Oak Decline (AOD) trees had greater macronutrient availability in root zone soils
- AOD trees had greater nutrient content in litter added to the root zone soil
- AOD root zone soils had altered microbial community structure & reduced CO_2_ emissions
- We conclude that AOD trees have poor physiological nutrient management and less active rhizosphere compared to healthy trees.

## 1. Introduction

Temperate woodlands and forests are important ecosystems that provide multiple ecosystem services such as climate regulation, carbon and nutrient cycling, air and water purification, habitat provision for biodiversity conservation, and recreational services for human wellbeing (de Gouvenain and Silander, 2017; Rawat et al., 2022; Silander, 2001). In the UK, expansion of woodlands is one of the strategic priorities to reach net zero carbon emissions (DEFRA, 2022; MacKenzie et al., 2024; MOJ, 2024). Similarly for biodiversity, the majority of the native species of conservation interest in the UK rely on forests for their survival (Quine and Humphrey, 2010). Despite the importance of forests in terms of ecosystem services and land-use policy for climate change mitigation, forests are threatened by several anthropogenic and environmental factors, such as pollution, climate change, forest degradation and fragmentation. However, one of the most important threats is decline in tree health, driven by pests and pathogens, which is further exasperated by other anthropogenic environmental degradation (Gilligan et al., 2013).

The current woodland cover of the UK is 2.84 million hectares, which is about 12% of the UK land area (MacKenzie et al., 2024). However, there are ambitious plans across the UK to drastically increase the woodland cover (DEFRA, 2022). English Oak (Quercus robur), also known as Pedunculate Oak is an important species for such expansion plans, considering that it is a native tree species that is known to provide great benefits for local biodiversity and very effective for climate change mitigation. English Oak is the second most common tree in the UK, and supports around 2300 species (Mitchell et al., 2019). English oak is also one of the most suitable species for efficient carbon sequestration. Recent Free Air Carbon Enrichment (FACE) experiments on mature oak trees have shown that those grown under elevated CO_2_ levels, as part of the CO_2_ fertilization effect, can absorb more CO_2_ from the atmosphere, aiding in climate change mitigation (MacKenzie et al., 2024). This contradicts observations in such FACE experiments on mature woodlands dominated by other species such as Eucalyptus, where elevated CO_2_ did not have any significant impact on tree uptake (Jiang et al., 2020), further emphasising the importance of oak species for climate change mitigation in temperate regions.

Tree health decline is a major threat to Oak, either through diseases caused by attack or infections of pests and pathogens that disrupt normal physiological functions, or by general decline caused by multiple environmental stressors and physiological factors that impair growth, resilience and overall health. In the UK, one such active infection classified as Acute Oak Decline (AOD) has become more and more prevalent over the recent years affecting woodlands in Southern and Eastern England (Denman et al., 2014; Denman and Webber, 2009). The disease is however less prevalent in Northern England and notably not identified in Scotland or Northern Ireland. This indicates the role of climate that relatively warmer, drier and draught prone weather in the south of England may support the disease. In fact, extreme weather conditions such as drought or water-logging, some geographical features such as lower elevation, proximity to industrial areas, and pollution especially nitrogen and sulphur deposition, were found to be abiotic indicators for the susceptibility and prevalence of oak decline in different European regions (Brown et al., 2018; Thomas et al., 2002). The AOD disease is characterised by multiple symptoms such as bleeding cankers (bleeding lesions on the stem at 1-2 m height, sometimes extending high into the canopy); deterioration in crown and canopy cover; and “D” shaped holes on the stem caused by Agrilus biguttatus beetle (Denman et al., 2014; Observatree, 2016; Reed et al., 2018). The presence of microbial pathogens (Brennaria goodwinii, Gibbsiella quercinecans & Lonsdalea Brittanica) and Agrilus beetle form an integral part of biotic indicators for susceptibility and prevalence of AOD (Brown et al., 2015; Gathercole et al., 2021). In fact, AOD is not prevalent in parts of the British Isles where Agrilus beetles are absent such as the island of Ireland. It should however be noted that AOD is prevalent in wider Continental Europe (Fernandes et al., 2022; González and Ciordia, 2020) and reported in places as far as Iran (Araeinejhad et al., 2024).

Acute oak decline is potentially caused by these biotic and abiotic factors, however as it is observed in different forests that just the presence of pathogens may not lead to decline symptoms in individual trees. Within a woodland, it is common to observe trees with acute decline symptoms surrounded by healthy oak trees with no sign of any decline. Some of the infected trees recover to full health, and some die within three years. COD in comparison is more passive general decline in health caused by multiple environmental stressors. Similar to AOD, COD is widely documented across the UK and Europe and is characterized by a gradual decline in tree health, marked by crown thinning, fine twig shedding, and dieback of scaffold branches, with failing root health thought to be a primary driver(Denman et al., 2022).

Both COD and AOD align with a decline spiral model, where environmental and biological predisposing, inciting, and contributing factors interact over time, exacerbating tree health deterioration (Denman et al., 2022; Gosling et al., 2024). However, it is not known what makes a particular tree within a woodland susceptible to decline in the presence of these biotic and abiotic conditions that favour AOD or COD. Hence soil, which is a source for all macro- and micro-nutrients required for healthy aboveground flora becomes a focus to understand the occurrence of oak decline symptoms (Powell et al., 2020). Soil is an integral part of any forest ecosystem, providing all the macronutrients and services that are essential for plant growth, aiding of cycling of elements including pollutants, and in carbon sequestration. Soil physics, chemistry and biology influence each other and form the basis of soil health (Powell et al., 2020). Soil microbial communities alongside soil physico-chemistry regulate the availability of different nutrients to plants (Brussaard et al., 2007). Soil microbial communities also comprises plant pathogens, alongside other beneficial microbes that may directly compete with the pathogens, or have antagonistic properties to pathogens, or develop disease resistance and resilience in plants (Hao and Ashley, 2021). For example, copper, cadmium and zinc are elements with known antimicrobial properties, and among the most common heavy metal contaminants in soils (Huang et al., 2019). So, it is essential to monitor availability of trace elements alongside nutrient availability and changes in microbial communities to understand the linkages between tree and soil health.

Forest litter plays an important role in nutrient cycling, and is the main source of nutrient addition to soil (Ge et al., 2013). In addition to litter chemistry influencing soil microbial communities and nutrient dynamics (Cleveland et al., 2007; Nemergut et al., 2010; Yang and Chen, 2009), it also provides information on pollution levels in the atmosphere (Lamano Ferreira et al., 2017), plant nutrient status and allocations, and several other indicators for ecosystem health and nutrient dynamics (Sayer et al., 2020; Sayer and Tanner, 2010). Some of the litter properties such as litter pH, phosphorus and nitrogen content, and carbon nitrogen ratio were known to influence soil nutrient availability for plants (Sayer et al., 2020; Tao et al., 2019; Yang and Chen, 2009). Alongside soil nutrient mineralisation, forest litter also forms the base for soil organic matter formation, and influences the carbon cycling and carbon balance of a forest ecosystem (Austin and Ballaré, 2010). Monitoring of leaf litter quantity and quality is essential in understanding changes in soil microbial communities, biogeochemical dynamics and their relation to tree health.

We monitored soil biogeochemical dynamics, microbial communities and above ground litter addition in root zone soil of oak trees of different health status (Acute Oak Decline, Chronic Oak Decline and Healthy) for a year (October 2022 to August 2023) in an oak dominated woodland. Our aim was to examine how soil properties and microbial communities vary with tree health and to explore potential feedback, via litter inputs, between soil biogeochemistry and tree decline. Our specific objectives were to

1. Identify the differences in soil biogeochemical properties, soil microbial communities, aboveground leaf litter properties between healthy and diseased trees covering temporal and soil depth scales.
2. Explore the complex relationship between tree health and soil biogeochemical dynamics and identify key effects of tree health on forest biogeochemical cycles.
3. Quantify the impact of tree health status on change in soil total carbon and nutrient content over the monitoring period of one year

## 2. Materials and Methods

### 2.1. Study site and sampling design

This study was carried out in Stoneymore Wood (Latitude <u>51.693396, Longitude 0.371544)</u>, which is part of the larger Writtle Forest woodland complex in Essex, England, UK. Stoneymore Wood is an ancient semi-natural woodland dominated by mature (> 75 years old) oak (Q. robur) with hornbeam (Carpinus betulus L.) as a secondary species (Booth, 2021). Oak decline has previously been documented in Writtle Forest (Booth, 2021) and the study focussed on selected oak trees within a 1 hectare area of the Stoneymore compartment where a cohort of 192 oak trees had been mapped in 2013 (Booth, 2021) and a census of above-ground decline symptoms taken in 2013, 2016, 2019 (Booth, 2021) and 2022.

For this study, three trees were selected for each of the three treatments (tree health status), making a total of nine experimental trees. The three treatments were 1) trees showing the main distinctive symptom (bleeding cankers) of Acute Oak Decline (AOD), 2) trees showing Chronic Oak Decline (COD) symptoms, and 3) Healthy (HEAL) trees with no signs of AOD or COD symptoms. The classification of health status was based on the longitudinal survey data of the mapped cohort and criteria for selection for each of the treatments is described in Figure 1. The map of Stoneymore wood and tree locations are given in the supplementary information (Supplementary Fig. 1). AOD trees had exhibited bleeding cankers since at least 2016, with fresh cankers (visible exudate) recorded in 2022. These trees had at ≥ 12 bleeding cankers (both recent and old) and an above-normal (15-45%) or excessive (45-85%) percentage of deadwood in the canopy, as classified based on crown transparency and the proportion of deadwood relative to the remaining crown. COD trees had no active bleeding cankers, though one tree had a small number (≤3) of inactive bleeds recorded in 2022. Healthy trees had no active, old, or callused bleeds and deadwood levels (10-15%) in the canopy were classified as ‘normal’. Further details of the symptom history of the studied trees are also given in the supplementary information (Supplementary Table 2).

**Figure 1:**
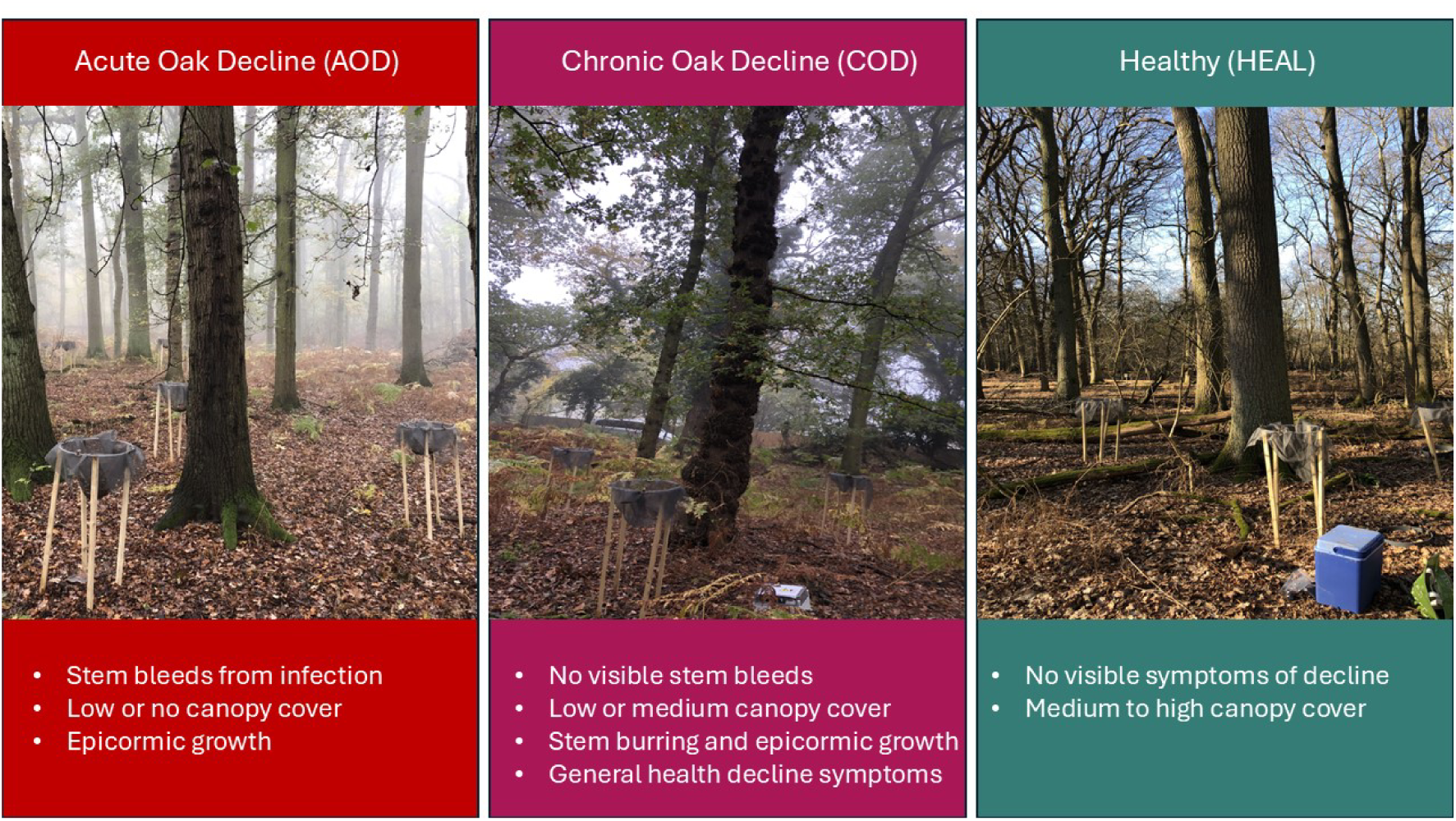
Classification of tree health status (treatments)

The soil from root zone of these trees was sampled once every four weeks from October 2022 to August 2023 (Exact dates of sampling are given in supplementary information-Supplementary Table 3). The sampling period covers all four seasons: Autumn (October-November 2022), Winter (December 2022 – February 2023), Spring (March – May 2023) and Summer (June – August 2023). The ‘regular’ monthly sampling was carried out, by collecting soil samples from 3 marked measurement points at locations within 1.5 m distance from the tree stem around each of the selected trees. The measurement points were marked by the placement of leaf litter traps (Fig. 1), and greenhouse gas (GHG) measurement chambers positioned near each of the three designated locations around each tree. Monthly soil sampling was carried out using graduated Bulb Planter, sampling to a depth of 10 cm. Within each season, one deep soil sampling was carried out using an Undisturbed Soil Corer from Eijelkamp (Giesbeek, Netherlands), sampling to a depth of 40 cm. The 40 cm core was separated into 10 cm segments and packed in airtight ziplock bags in the field. The soil samples were then carried in a cool box to the labs in The University of Reading immediately after the sampling. The soil samples were then sieved field moist using 2 mm sieve for 0-10 cm layers, and 4 mm sieve for deeper layers. A portion of each sample was used for KCl extraction immediately after sieving, while another portion was air dried in a drying cabinet for Mehlich 3 extraction. Additionally, a portion of fresh sieved sample was frozen at -20°C and freeze-dried for phospholipid fatty acid (PLFA) analyses. Finally, another portion was oven dried (105 °C) to determine moisture content, and to conduct further chemical analyses including total carbon and nutrients. For PLFA analyses, the samples from three sampling locations around each tree were pooled together, making it one composite sample from each tree. Methods for soil physico-chemical (section 2.2) and microbial (section 2.3) analysis are further described below.

Leaf litter samples were also collected monthly using self-constructed leaf litter traps. The litter traps were constructed based on the instructions from the manual produced by International Co-operative Programme on Assessment and Monitoring of Air Pollution Effects on Forests (ICP Forests) (Ukonmaanaho et al., 2020) using 50 cm diameter metal wire rings (Dennels, UK) as frames, mounted on wooden stakes of 120 cm length (Suregreen, UK) that were hammered into the soil.

Water- and fire-resistant insect mesh was clipped onto the metal rings, and used as the net to trap litter. The leaf litter samples were collected every four weeks at the same time as soil sample collection, packed in ziplock bags and transported to the labs on the same day for processing and analyses. For leaf litter chemistry analyses the samples from 3 positions for each tree were pooled together to form one composite sample for each individual tree. GHG Samples were collected every four weeks from the headspace of static chambers, using a syringe and glass exetainer vials (refer to section 2.5), and transported to the lab for analyses.

### 2.2. Soil physico-chemical analyses

#### 2.2.1 Soil temperature and moisture

The procedures used for soil analyses were based on Dhandapani et al. (2023b); Dhandapani et al. (2022). Soil temperature was measured in situ, using a digital thermometer RS-40 made by RS instruments (Corby, UK). For gravimetric moisture, 10 g of fresh soil was dried in an oven at 105° C for 24 hours. Then gravimetric moisture was calculated based on mass of water lost on oven drying. Volumetric moisture was measured in situ, using a digital volumetric moisture meter, ThetaProbe® (Delta-T devices, Cambridge, UK).

#### 2.2.2 pH and electrical conductivity

For pH and electric conductivity measurements, a 5 mL volume of soil sample was diluted in 10 mL deionised water and shaken for 30 mins at 25 rpm in a rotary shaker. The pH and electrical conductivity were measured using HI 9812-5 portable meter made by Hanna Instruments (Leighton Buzzard, UK).

#### 2.2.3 Plant available nutrients and trace elements in soil

Inorganic nitrogen (N) in the forms of Nitrate-N and ammonium-N concentrations were determined using potassium chloride (KCl) extraction. For this, 50 mL 1M KCl solution was added to 10 g of fresh 2-mm sieved soil (Buurman et al., 1996). Samples were shaken for 1 hour with an orbital shaker, then filtered through Whatman GF/A 15 cm filter paper, and finally analysed using Skalar San++ elemental analyser (Skalar, Breda, Netherlands).

Other available nutrients and trace elements were measured using Mehlich-3 extraction. For this, 2 g of 2mm sieved air-dried soil samples were treated with 20 mL Mehlich-3 solution (0.2 M CH_3_COOH, 0.25 M NH_4_NO3, 0.015 M NH_4_F, 0.013 M HNO_3_, and 0.001 M EDTA, at pH 2.45) (Bolland et al., 2003). Samples were shaken for 5 minutes at 40 rpm in a rotary shaker. The samples were then filtered through Whatman no. 42 filter paper, diluted and finally analysed using inductively coupled plasma optical emission spectroscopy (ICP-OES; Avio 550 Max Perkin Elmer, Beaconsfield, UK).

#### 2.2.4 Total Carbon and Nutrients in Soil

For analysing total carbon (C) and nitrogen (N) content, all samples were oven dried (105°C for 24 h) and finely ground using a Fritsch Pulverisette 5 ball mill (Brackley, UK). Samples (10 mg) were then weighed in tin cups which were folded and sealed prior to analysis by Flash 2000 Elemental Analyser (Thermo Fisher Scientific, Oxford, UK).

Soil total nutrient concentrations were analysed using ICP-OES following sample digestion. For this, approximately 0.5 g of oven dried (105°C for 24 h) and ground soil were weighed in microwave digestion tubes (MARSXpress 6 vessels, CEM Microwave Technology Ltd., Buckingham, UK) and 9 mL of nitric acid + 3 mL of hydrochloric acid was added to each sample. The digestion tubes were then placed in a MARSXpress microwave (CEM Microwave Technology Ltd., Buckingham, UK.) and run at 175°C with a ramp up time of 5 min 30 sec, followed by a hold time of 4 min 30 sec at 175°C. The digested samples were then filtered and diluted using milliQ water, and were then analysed using Avio 550 Max ICP-OES.

### 2.3. Soil microbial community structure

Microbial community phenotypic structure was determined by phospholipid fatty acid (PLFA) analysis as described in Dhandapani et al. (2020). PLFAs were extracted from 5 g freeze-dried soil samples using a modification of the method described by Frostegard et al. (1991). Lipids were extracted using Bligh & Dyer extraction method (Bligh & Dyer, 1959). The extracted lipids were then separated into neutral, glycol and polar lipid (containing phospholipids) fractions using Megabond Elut® silica gel columns. The extracted polar lipids were then methylated by mild alkaline methanolysis and converted into fatty acid methyl esters.

The dried fatty acid methyl esters were suspended in 200 µl of hexane. The samples were then analysed using an Agilent 6890N Gas Chromatography Flame Ionisation Detector (GC-FID) instrument (Agilent Technologies, Didcot, UK). One μl of each sample was injected into the GC in split-less mode. The carrier gas was helium with the constant pressure of 18psi. The initial oven temperature in GC was 60°C; this was maintained for 1 min and then programmed to 250°C at the rate of 2.5°C min^-1^. The constant temperature of 250°C was maintained throughout the run. The results were displayed as a chromatogram of retention times of the compounds.

The fatty acids i15:0, a15:0, i16:0, i17:0 and a17:0 were classified as Gram-positive biomarkers (Wilkinson et al., 2002). Methyl-branched fatty acids 10me16:0 and 10me18:0 were described as the biomarkers for actinobacteria (Wilkinson et al., 2002; Moore-Kucera & Dick, 2008), a group that belongs to Gram-positive bacteria. The relative abundance of Gram-negative bacteria was calculated using 2-OH 14:0,3 OH 14:0, 2 OH 16:0 16:1n11t, 16:1, 16:1n7, 16:1n7t, cyc17:0, and 18:1n9t as biomarkers (Wilkinson et al., 2002; Kaiser et al., 2010). 18:2n6 and 18:1n9 were used as fungal biomarkers (Vestal & White, 1989; Wilkinson et al., 2002; Kaiser et al., 2010). Fatty acids such as 14:0, 15:0, 16:0, 17:0, 2OH 16:0, 18:0, Cyc19:0 and 20:0 were classified as general bacterial fatty acids (Wilkinson et al., 2002). Fatty acids 20:5n6 and 20:5n3 were classed as Micro Eukaryotes. Fatty acids such as 17branched, 19:1n6 and 19:1n8 could not be classified into any specific groups. The ratio of Cyclopropane fatty acids (cyc17:0) to their monoenoic precursor (16:1n7) and the ratio of total saturated fatty acids (14:0, 15:0, 16:0,17:0, 18:0, 20:0) to mono-unsaturated fatty acids (16:1, 16:1n11t, 16:1n7, 16:1n7t, 16:1n5 a17:1n10, 18:1n9, 18:1n9t, 18:1n10, 19:1n6, 19:1n8, 20:1n9) were used as indicators of stress (such as reduced carbon and nutrient availability) and other ecological conditions such as flooding (Bossio & Scow, 1998).

### 2.4. Litter analyses

#### 2.4.1 Sample preparation

The collected litter was oven dried in paper bags at 70° C for 72 hours. The dried samples were sorted into leaves, wood, seeds and flowers, and dry weight was recorded. The dried and separated litter components were then finely ground using Fritsch Pulvereisette 14 classic line rotor mill (Brackley, UK).

#### 2.4.2 Litter chemistry

For pH measurements, 5 mL volume of ground litter sample was diluted in 10 mL deionised water and shaken for 30 mins at 25 rpm in a rotary shaker. The pH was measured using accumet® AE150 pH meter made by Fisher Scientific (Loughborough, UK).

For analysing total C and N content, finely ground samples were weighed to 5 mg in tin cups, folded, sealed and analysed in Flash 2000 Elemental Analyser (Thermo Fisher Scientific, Oxford, UK).

Leaf litter total nutrient concentrations were analysed using ICP-OES following sample digestion. For this, approximately 0.5g of oven dried (70°C for 72 h) and ground litter samples were weighed in microwave digestion tubes (MARSXpress vessels, CEM Microwave Technology Ltd., Buckingham, UK), and 8 mL of nitric acid was added to each sample. The digestion tubes were then placed in a MARSXpress microwave (CEM Microwave Technology Ltd., Buckingham, UK.) and heated to 200°C with a ramp-up time of 20–25 minutes. The temperature was then maintained at 200°C for 10 minutes. The digested samples were then filtered, diluted using milliQ water, and were then analysed using Avio 550 Max ICP-OES.

### 2.5. Greenhouse gas analyses

CO_2_, CH_4_ and N_2_O emissions from the soil surface were measured using static chambers, based on the guidelines from Clough et al. (2020). The closed chambers used were opaque with a height of 15 cm and an inner diameter of 40 cm. Collars of 15 cm height were installed in the site. The collars were inserted to a depth of 10 cm, however in some of the sampling locations the insertion depth was variable because of the logistical difficulties of inserting to a depth of 10 cm in forest soils.

Nevertheless, the collar height was measured in 5 different locations, and the mean value for each of the collar was used for the calculation of height of each chamber and subsequent gas flux calculations. The chambers and collars were manufactured in-house using 40 cm diameter thick PVC pipes. During each measurement the chamber was carefully placed on the collar, and a gasket was used to provide a gas tight seal. Gas samples were taken at 0 min and at 60 min using a gas tight syringe to take a sample from the chamber headspace via a gas sampling port fitted to the top of the chamber. Gas samples (15 ml) were injected to pre-evacuated exetainer vials (12 ml) creating over-pressure. The samples were then sent to University of Nottingham for analyses using a gas chromatograph equipped with electron capture (N_2_O), flame ionisation (CH_4_) and thermal conductivity (CO_2_) detection, which provided gas concentrations in ppm. The gas concentrations in ppm were converted to mg m^-2^ hr^-1^ (for CO_2_ ) and µg m^-2^ hr^-1^ (for CH_4_ and N_2_O) as described in Dhandapani et al. (2022)) using the Ideal Gas Law: PV=nRT , where: P = atmospheric pressure; V = volume of headspace; n = number of moles (mol); R = universal Gas Constant law (8.314 J K^-1^mol^-1^) and T = temperature in kelvin (K), with the conversion factor, 1 mol of CO_2_ = 44.01 g, 1 mol CH_4_ = 16.02 g and 1 mol of N_2_O is 44.012 g. The change in gas concentration within the chamber volume (cm^3^) for soil surface area (m^2^) covered were fitted using linear regression.

### 2.6. Statistical analyses

#### 2.6.1 Soil Chemistry

All statistical analyses were carried out using Genstat® 23^rd^ edition (VSN international, Hemel Hempstead). Linear mixed models with Restricted Maximum Likelihood (REML) were used to determine the difference between treatments for soil chemistry. Surface soil (0-10 cm) chemistry parameters were modelled as dependent variables using a linear mixed model, with interactions between tree health treatment (n=3) and measurement period (n=12) as categorical fixed effects, and sampling positions (n=3 per tree) nested within individual trees used as random effects.

For the analysis of soil chemistry parameters including the effect of soil depth (0-10, 10-20, 20-30 and 30-40 cm), linear mixed models were run with interactions between tree health treatment (n=3) and depth (n=4) as categorical fixed effects, and sampling positions (n=3 per tree) nested within individual trees used as random effects.

To determine the changes in total carbon and nutrient contents from the start of the monitoring period to the end of the monitoring period, linear mixed models were used, with total carbon and nutrient contents as dependent variables, and treatment (n=3), measurement period (n=2) and soil depth (n=4) as categorical fixed effects, with sampling positions (n=3 per tree) nested within individual trees used as random effect.

#### 2.6.2 Soil Microbial Community Structure

The significance of differences between treatments for relative abundance of different microbial groups were evaluated using two-way analysis of variance (ANOVA), with interactions between treatment and season as fixed effects. Mixed models could not be used for PLFA data, because of the pooling of field samples for each tree, making the data not suitable for mixed models. Principal component analysis (PCA) was performed for the dataset containing relative abundance of all the individual fatty acids, to identify the main axes of variance with soil microbial PLFAs, existing patterns between measured parameters, and to measure the difference between different treatments and seasons.

#### 2.6.3 Soil Greenhouse Gas emissions

Linear mixed models with Restricted Maximum Likelihood (REML) were used to determine the difference between treatments in GHG emissions. Different GHG (N_2_O, CH_4_ and CO_2_) were modelled as dependent variables using linear mixed model, with interactions between treatment (n=3) and measurement period (n=12) as categorical fixed effects, and sampling positions (n=3 per tree) nested within individual trees used as random effect.

#### 2.6.4 Litter Quantity and Quality

Linear mixed models with Restricted Maximum Likelihood (REML) were used to determine the difference between quantity of leaf litter. Weight of different components of leaf litter (leaf, wood, seed and flower) were modelled as dependent variables using linear mixed model, with interactions between treatment (n=3) and measurement period (n=12) as categorical fixed effects, and sampling positions (n=3 per tree) nested within individual trees used as random effect.

The significance of differences between treatments for litter chemistry parameters of different components of leaf litter were evaluated using two-way analysis of variance (ANOVA), with interactions between treatment and season as fixed effect. Mixed models could not be used for litter chemistry data, because of the pooling of field samples for each tree, making the data not suitable for mixed models.

## 3. Results

### 3.1 Surface soil monthly geochemical dynamics

Surface soil chemistry (0–10 cm) exhibited substantial temporal variation, while the effects of tree health status were relatively minor in comparison to this temporal variation. The monthly surface soil temperature ranged from the highs of 17.2°C in summer to the lows of 4.2°C in winter, and it was the only measured parameter that showed significant main effect of health status, however the variations were minimal (Figure 1; Table 1). Several other soil chemistry parameters such as pH, available forms of macronutrients such as Phosphorus (P), Sulphur (S) and potassium (K), available Aluminium (Al) showed significant interactions between treatment and measurement period. The general trend for these macronutrients and Al is that they showed higher availability in AOD root zone soil for most of the months except in summer months (especially August), where nutrient and Al availability under AOD dropped below COD and HEAL treatments, resulting in significant statistical interactions. Similarly soil electrical conductivity was particularly high for COD in the August measurement period driving significant interactions. Soil pH remained the lowest for AOD trees for most of the months except for the summer period, resulting in significant statistical interactions between treatment and measurement period.

**Table 1:** Linear mixed model (REML) for soil chemistry and nutrient availability, showing statistical significance of the effects of treatment (tree health status), sampling month, and the interactions between treatment and month. Statistically significant figures are presented in bold italics.

|  | Treatment | Month | Treatment*Month |
| --- | --- | --- | --- |
| Temperature | $F_{(2,6)}=7.06, p=0.027$ | $F_{(10,240)}=193.36, p<0.001$ | $F_{(20,240)}=1.04, p=0.415$ |
| Volumetric Moisture | $F_{(2,6)}=1.33, p=0.333$ | $F_{(10,240)}=70.06, p<0.001$ | $F_{(20,240)}=1.31, p=0.170$ |
| Gravimetric Moisture | $F_{(2,6)}=1.07, p=0.4$ | $F_{(11,264)}=28.37, p<0.001$ | $F_{(20,264)}=1.27, p=0.193$ |
| pH | $F_{(2,6)}=0.38, p=0.697$ | $F_{(11,263.1)}=10.81, p<0.001$ | $F_{(20,240)}=1.79, p=0.018$ |
| Electrical Conductivity | $F_{(2,6)}=0.02, p=0.979$ | $F_{(11,263.1)}=42.45, p<0.001$ | $F_{(11,263.1)}=2.48, p<0.001$ |
| Macronutrients |  |  |  |
| log Ammonium-N | $F_{(2,6)}=0.73, p=0.521$ | $F_{(11,264)}=2.23, p=0.014$ | $F_{(22,264)}=0.78, p=0.752$ |
| logNitrate-N | $F_{(2,4.5)}=0.93, p=0.459$ | $F_{(11,60.2)}=3.26, p=0.002$ | $F_{(18,59.6)}=0.83, p=0.66$ |
| P | $F_{(2,6)}=3.81, p=0.085$ | $F_{(10,239.1)}=14.77, p<0.001$ | $F_{(20,239.1)}=1.65, p=0.042$ |
| Mg | $F_{(2,6)}=0.03, p=0.967$ | $F_{(10,239.1)}=10.02, p<0.001$ | $F_{(20,239.1)}=0.55, p=0.942$ |
| logS | $F_{(2,6)}=1.01, p=0.419$ | $F_{(10,239.1)}=31.33, p<0.001$ | $F_{(20,239.1)}=2.02, p=0.007$ |
| logCa | $F_{(2,6)}=0.09, p=0.915$ | $F_{(10,239)}=6.63, p<0.001$ | $F_{(20,239)}=0.34, p=0.997$ |
| K | $F_{(2,6)}=0.78, p=0.498$ | $F_{(10,239)}=15.71, p<0.001$ | $F_{(20,239.1)}=2.05, p=0.06$ |
| Micronutrients |  |  |  |
| Fe | $F_{(2,6)}=1.12, p=0.386$ | $F_{(10,239.1)}=22.68, p<0.001$ | $F_{(20,239)}=1.57, p=0.062$ |
| logMn | $F_{(2,6)}=1.4, p=0.317$ | $F_{(10,239.1)}=7.78, p<0.001$ | $F_{(20,239)}=0.66, p=0.865$ |
| logZn | $F_{(2,6)}=0.63, p=0.565$ | $F_{(10,239)}=11.48, p<0.001$ | $F_{(20,239)}=0.57, p=0.928$ |
| logCu | $F_{(2,6)}=0.48, p=0.639$ | $F_{(10,239.1)}=28.57, p<0.001$ | $F_{(20,239.1)}=0.81, p=0.701$ |
| logNi | $F_{(2,6)}=0.37, p=0.705$ | $F_{(10,239)}=17.5, p<0.001$ | $F_{(20,239)}=0.96, p=0.511$ |
| logNa | $F_{(2,6)}=0.13, p=0.879$ | $F_{(10,239)}=24.38, p<0.001$ | $F_{(20,239)}=0.73, p=0.797$ |

Several macro and micro-nutrients such as nitrate-N, ammonium-N, magnesium (Mg), calcium (Ca), iron (Fe), zinc (Zn), nickel (Ni), manganese (Mn), copper (Cu), sodium (Na), alongside moisture showed significant measurement period effect, showing high temporal variation in soil nutrient availability. The general trend was that the nutrient availability was significantly higher in autumn for most nutrients (nitrate-N, ammonium-N, Ca, Mn and Zn) and in spring for certain nutrients (such as Cu), driving significant temporal variations in surface soil nutrient availability. Fe showed high fluctuations in availability throughout the year without any specific seasonal pattern.

### 3.2 Deep soil seasonal geochemical dynamics

Seasonal deeper soil sampling and analyses showed a strong depth effect and a clear treatment effect with increased nutrient availability in the AOD root zone soil. Nitrate-N, Ni and Na showed significant treatment effect, while ammonium-N, P, Mg and S showed significant interactions between treatment and season, emphasising the influence of tree health status on seasonal deep soil chemistry. All these nutrient concentrations were significantly greater in the surface layers of the AOD oak root zone, and then decreased with depth and reached similar levels as other treatments for nitrate-N, ammonium-N and phosphorus; and showed different trends of change with depth for Mg, Ni , Na and S. Notably S concentrations decreased with depth for AOD and HEAL, while COD showed increase with depth with greater S concentrations in deeper layers than what is observed in AOD root zone. Similarly, Mg and Ni concentrations in bottom two layers of HEAL root zone soil were greater than those of the other two treatments, even though HEAL followed similar trend of decrease as other two treatments at 10-20 cm depth. Na concentrations increased with depth for all three treatments however degree of change was different between the treatments at each depth level.

All the measured nutrients, alongside gravimetric moisture, electrical conductivity and pH also showed significant variation with depth. Moisture, electrical conductivity and most nutrient availability (except Mg, Na, S and Al) decreased with depth (Figure 4, 5 and Table 2). Al concentrations increased from surface to 10-20 cm and stayed at that higher level in deeper layers (Figure 4).

**Table 2:**
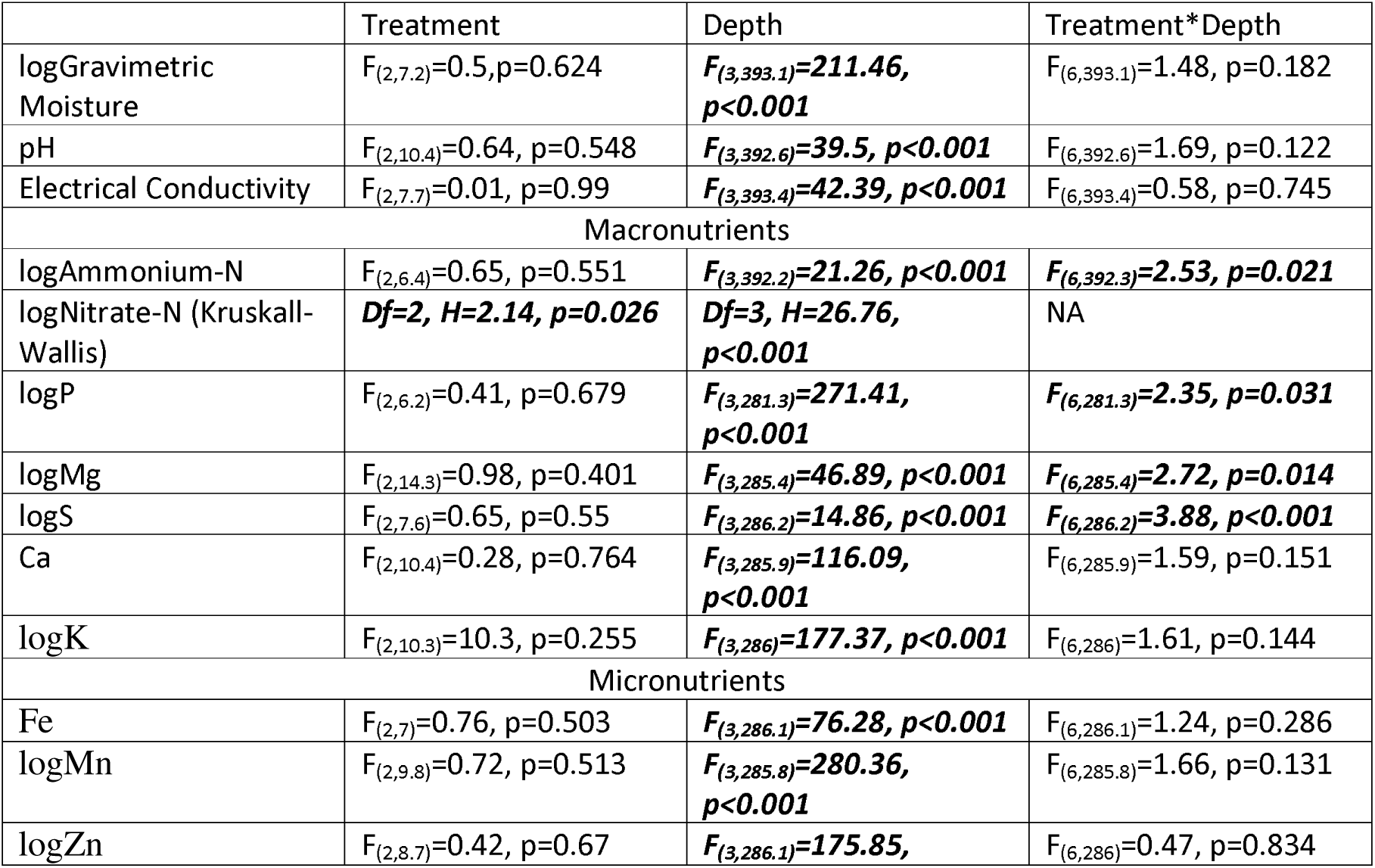
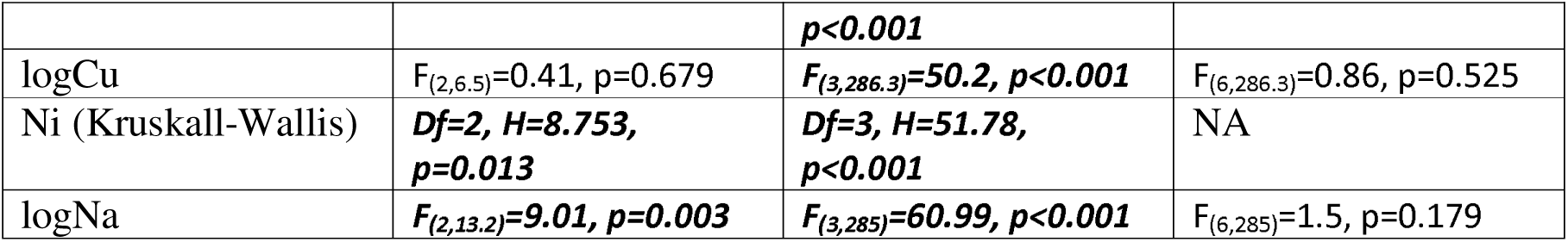
Linear mixed model (REML) for soil chemistry and nutrient availability, showing statistical significance of the effects of treatment (tree health status), depth, and the interactions between treatment and depth. Statistically significant figures are presented in bold italics.

**Figure 2:**
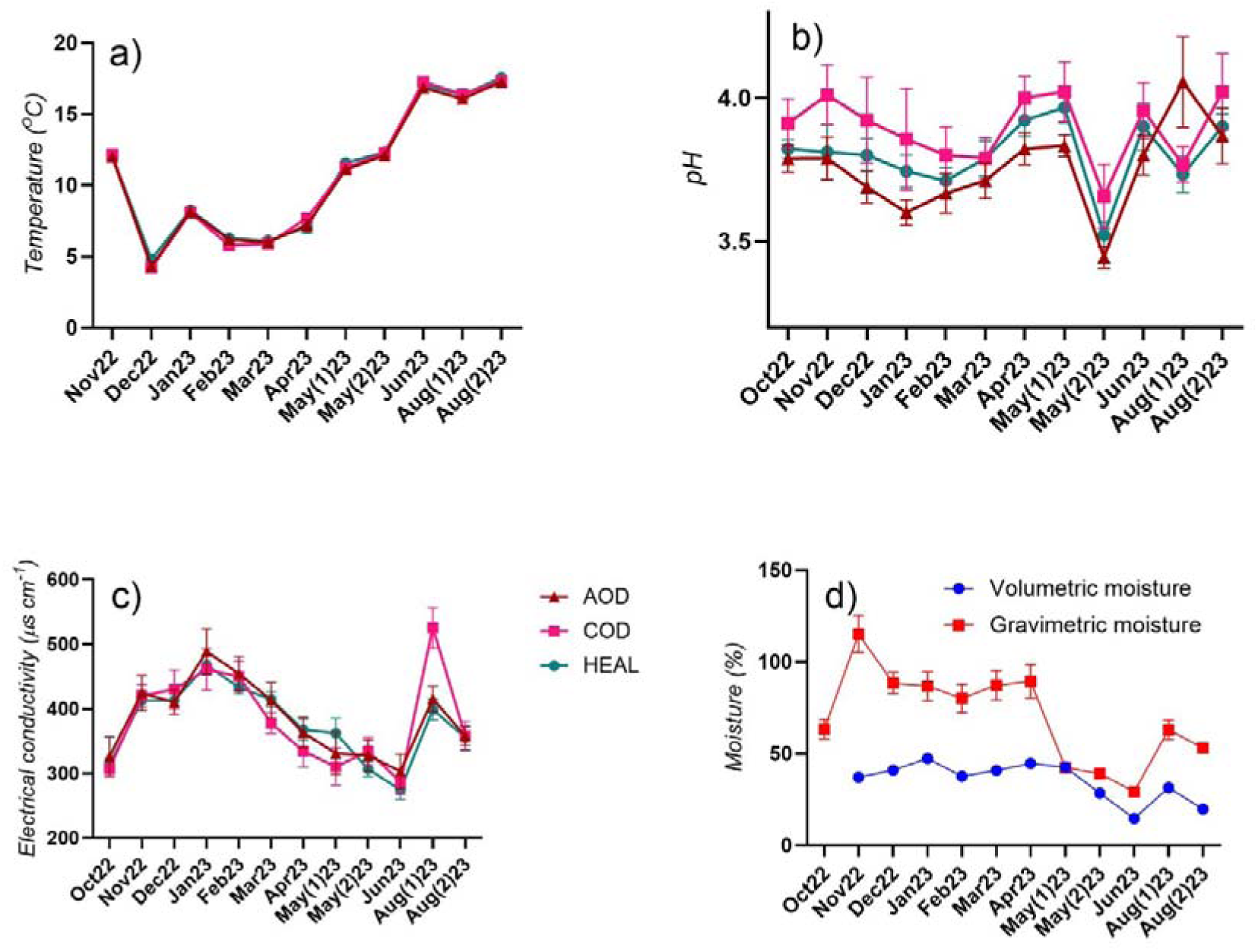
Effects of Treatment and Measurement period upon a) soil temperature, b) soil pH, c) soil electrical conductivity, and d) soil moisture. Only significant effects from linear mixed model (REML) are presented in the figure. Whiskers denote standard errors of means. Data points for which the error bars are not visible, the error bars are smaller than the symbols used for data points.

**Figure 3:**
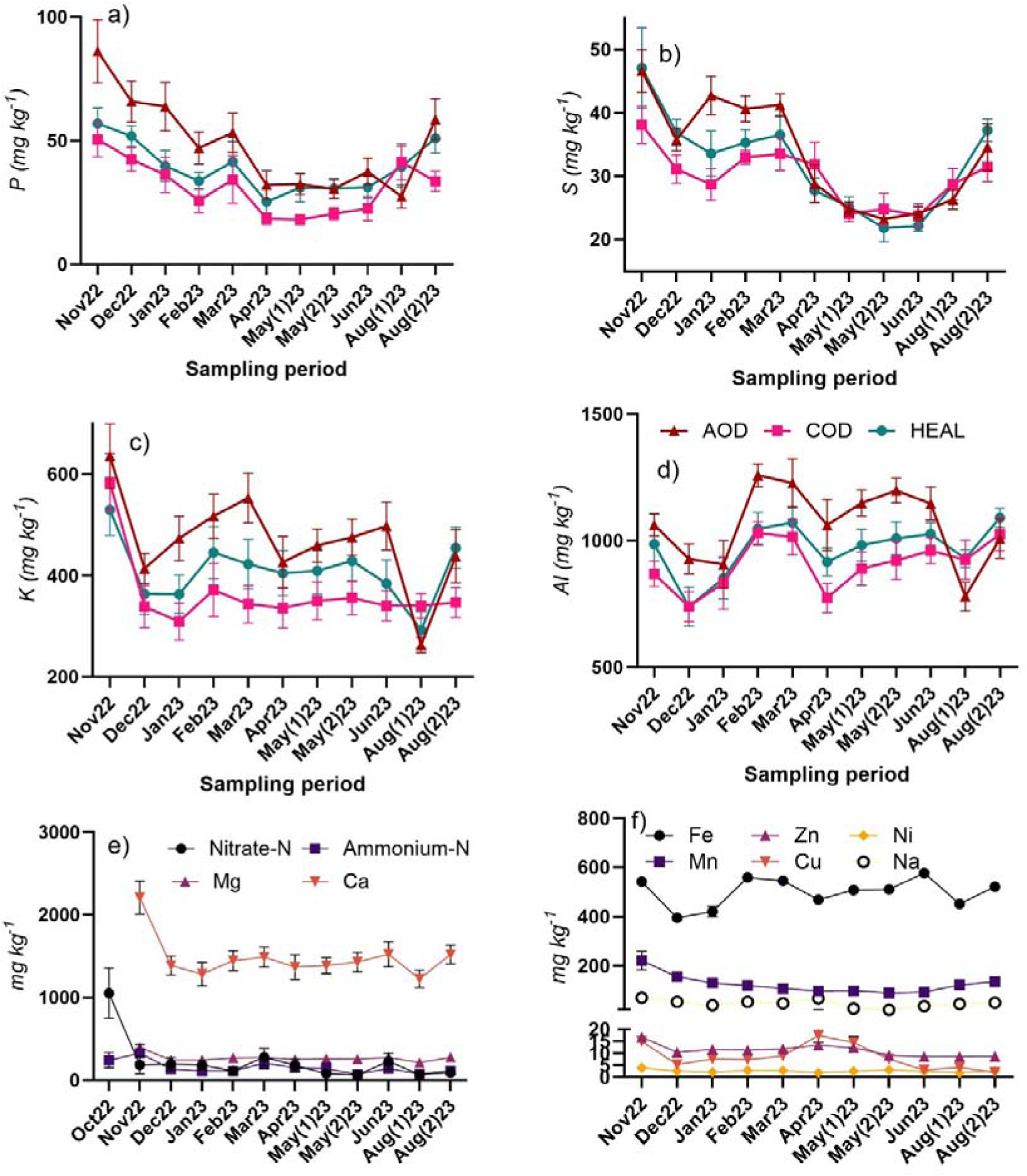
Effect of treatment and measurement period upon available (Mehlich 3 extractable) a) Phosphorus (P), b) Sulphur (S), c) Potassium (K), d) Aluminium (Al), e) other macronutrients (nitrate-N, ammonium-N, Mg and Ca), and d) micronutrients (Fe, Zn, Ni, Mn, Cu and Na). Only significant effects from linear mixed model (REML) are presented in the figure. Whiskers denote standard errors of means. Data points for which the error bars are not visible, the error bars are smaller than the symbols used for data points.

**Figure 4:**
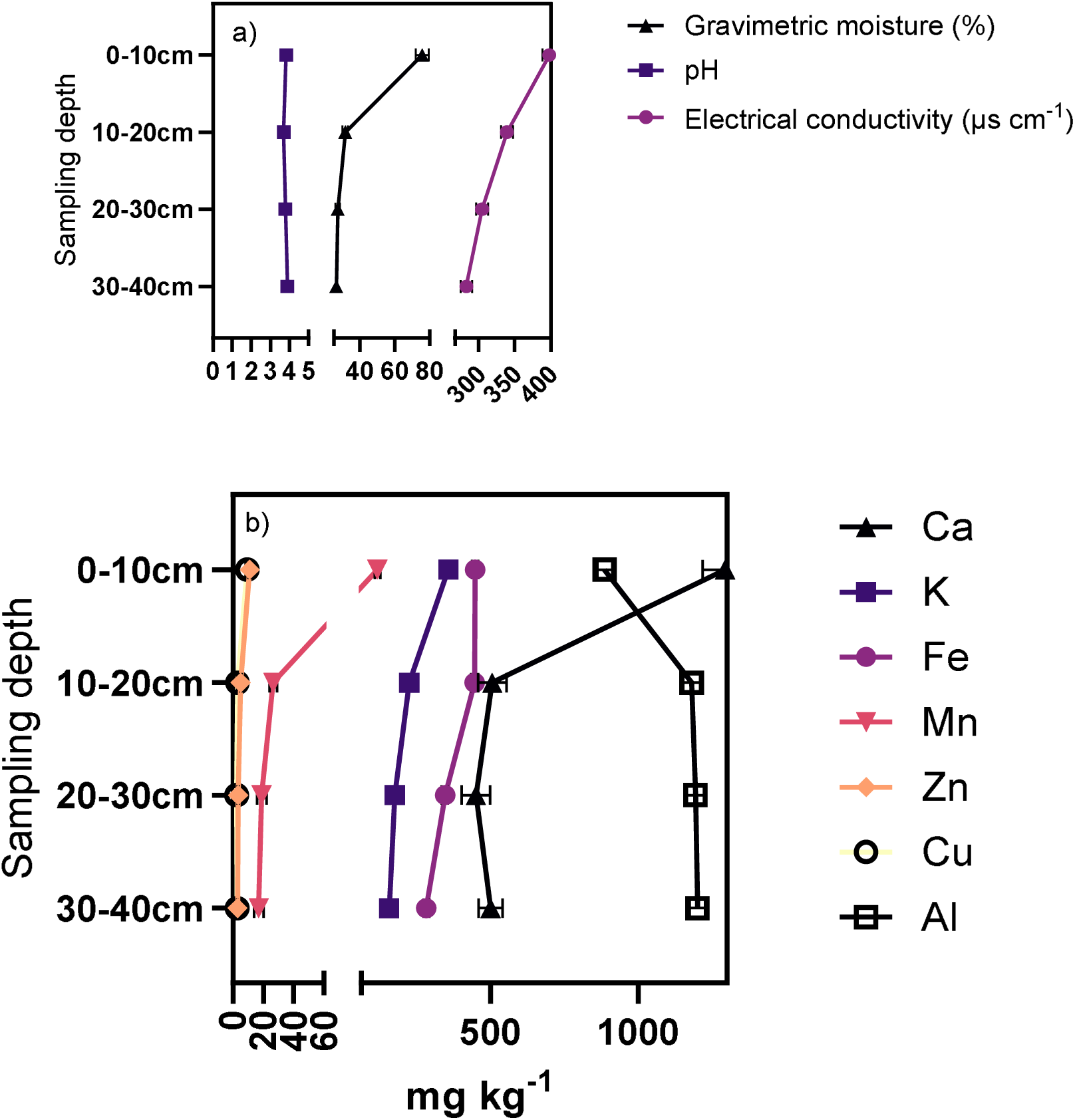
Effect of treatment and measurement period upon a) Soil chemistry parameters (gravimetric moisture, pH, electrical conductivity), b) available (Mehlich 3 extractable) macro- and micro-nutrients (Ca, K, Fe, Mn, Zn, Cu, Al). Only significant effects from linear mixed model (REML) are presented in the figure. Whiskers denote standard errors of means. Data points for which the error bars are not visible, the error bars are smaller than the symbols used for data points.

**Figure 5:**
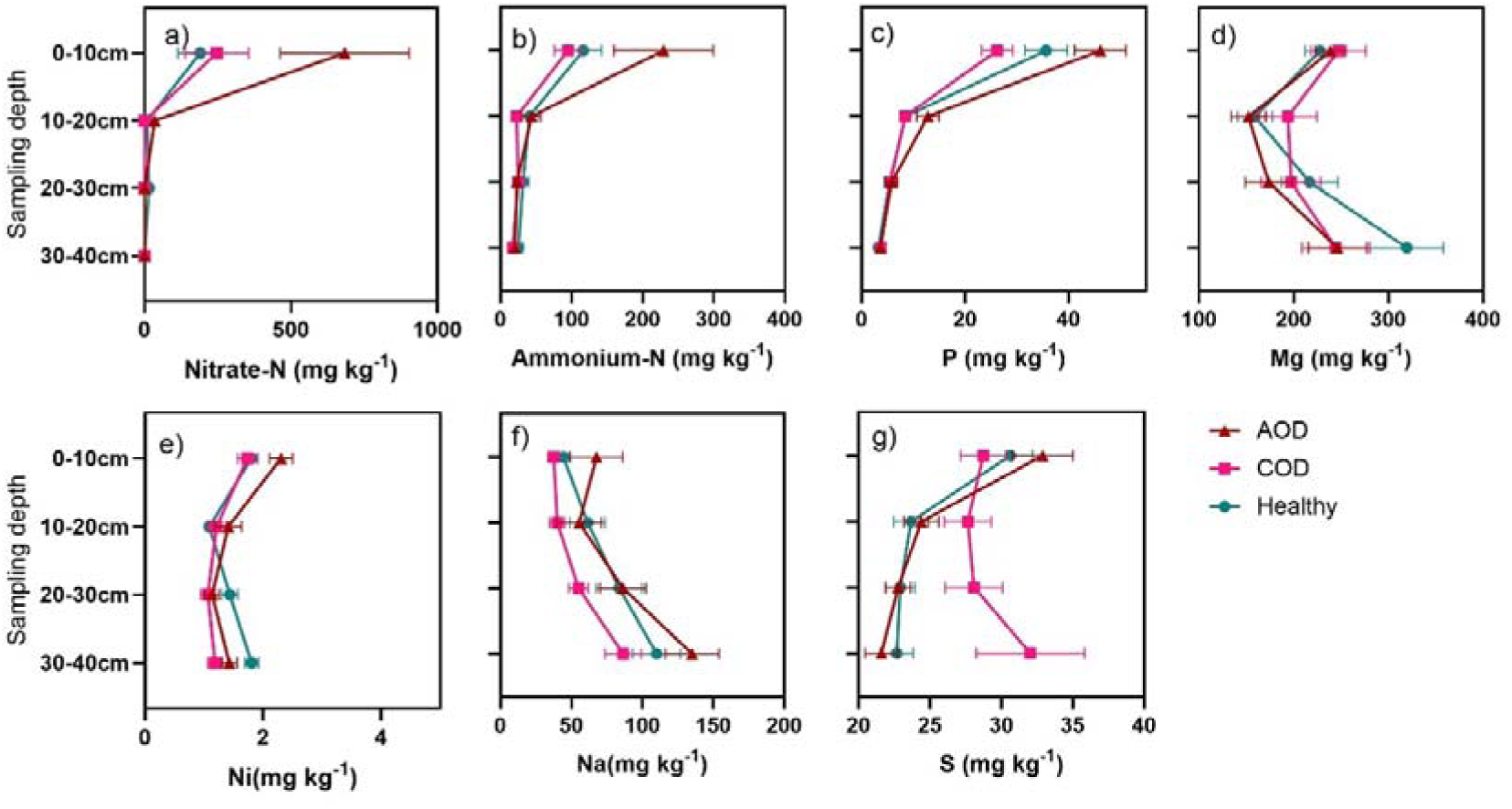
Effect of treatment and measurement period upon a) Nitrate-N, b) Ammonium-N, c) available P, d) Mg, e) Ni, f) Na and g) S. Only significant effects from linear mixed model (REML) are presented in the figure. Whiskers denote standard errors of means. Data points for which the error bars are not visible, the error bars are smaller than the symbols used for data points.

### 3.3 Annual change in total soil C and nutrients

There was no treatment effect observed on changes in total carbon and nutrient content in soils from the start of the monitoring period to the end of the monitoring period. However, there was a general decline in carbon and nutrient (N, P, Mg, K, Zn, Cu, Ni and Na) contents throughout the depth of the root zone soil (Figure 6, Table 3). Zn was the only element which showed slight yet significant increase in concentration at the end of monitoring period. Other nutrients showed significant changes with depth, with no increase or decrease during the monitoring period (Table 3). Carbon to nitrogen (C:N) ratio increased over the monitoring period, despite both carbon and nitrogen content showing similar trend of decrease over this period, pointing at the greater degree of loss of nitrogen than carbon.

**Table 3:** Linear mixed model (REML) for soil chemistry and total nutrient contents, showing statistical significance of the effects of treatment (tree health status), measurement period and depth. Statistically significant figures are presented in bold italics.

|  | Treatment | Measurement period | Depth |
| --- | --- | --- | --- |
| logC | $F_{(2,6)}=0.85, p=0.474$ | ***$F_{(1,183.1)}=4.97, p=0.027$*** | ***$F_{(3,183.1)}=118.15, p<0.001$*** |
| logN | $F_{(2,10.4)}=0.64, p=0.548$ | ***$F_{(3,392.6)}=39.5, p<0.001$*** | $F_{(6,392.6)}=1.69, p=0.122$ |
| C:N (Kruskal Wallis) | Df=2, H=5.002, $\chi^2=0.082$ | ***Df=1, H=17.75, $\chi^2<0.001$*** | ***Df=3, H=56.68, $\chi^2<0.001$*** |
| Other Macronutrients |  |  |  |
| P | $F_{(2,6)}=0.18, p=0.837$ | ***$F_{(1,82.2)}=12.62, p<0.001$*** | ***$F_{(3,182.2)}=137.92, p<0.001$*** |
| Mg | $F_{(2,6)}=0.11, p=0.895$ | ***$F_{(1,182.3)}=8.64, p=0.004$*** | ***$F_{(3,182.3)}=8.34, p<0.001$*** |
| logS | $F_{(2,6)}=0.29, p=0.755$ | $F_{(1,182.4)}=1.13, p=0.289$ | ***$F_{(3,182.4)}=193.63, p<0.001$*** |
| Ca | $F_{(2,6)}=0.14, p=0.871$ | $F_{(1,182.4)}=0.02, p=0.886$ | ***$F_{(3,182.4)}=129.25, p<0.001$*** |
| K | $F_{(2,6)}=0.01, p=0.985$ | ***$F_{(1,182.3)}=39.58, p<0.001$*** | $F_{(3,182.3)}=2.06, p=0.107$ |
| Micronutrients |  |  |  |
| Fe | $F_{(2,6)}=0.16, p=0.859$ | $F_{(1,182.4)}=0.03, p=0.858$ | ***$F_{(3,182.3)}=19.91, p<0.001$*** |
| logMn | $F_{(2,6)}=0.48, p=0.639$ | $F_{(1,182.1)}=0.07, p=0.787$ | ***$F_{(3,182.1)}=54.63, p<0.001$*** |
| logZn | $F_{(2,5.9)}=0.89, p=0.458$ | ***$F_{(1,171.2)}=13.64, p<0.001$*** | $F_{(3,169.9)}=0.4, p=0.75$ |
| Cu | $F_{(2,6)}=0.16, p=0.854$ | ***$F_{(3,183)}=44.2, p<0.001$*** | ***$F_{(3,183)}=25.95, p<0.001$*** |
| Ni | $F_{(2,6)}=0.05, p=0.955$ | ***$F_{(3,182)}=15.73, p<0.001$*** | ***$F_{(3,182.2)}=5.29, p=0.002$*** |
| logNa | $F_{(2,6)}=0.15, p=0.864$ | ***$F_{(1,182.2)}=11.64, p<0.001$*** | ***$F_{(3,182.2)}=11.64, p<0.001$*** |

**Figure 6:**
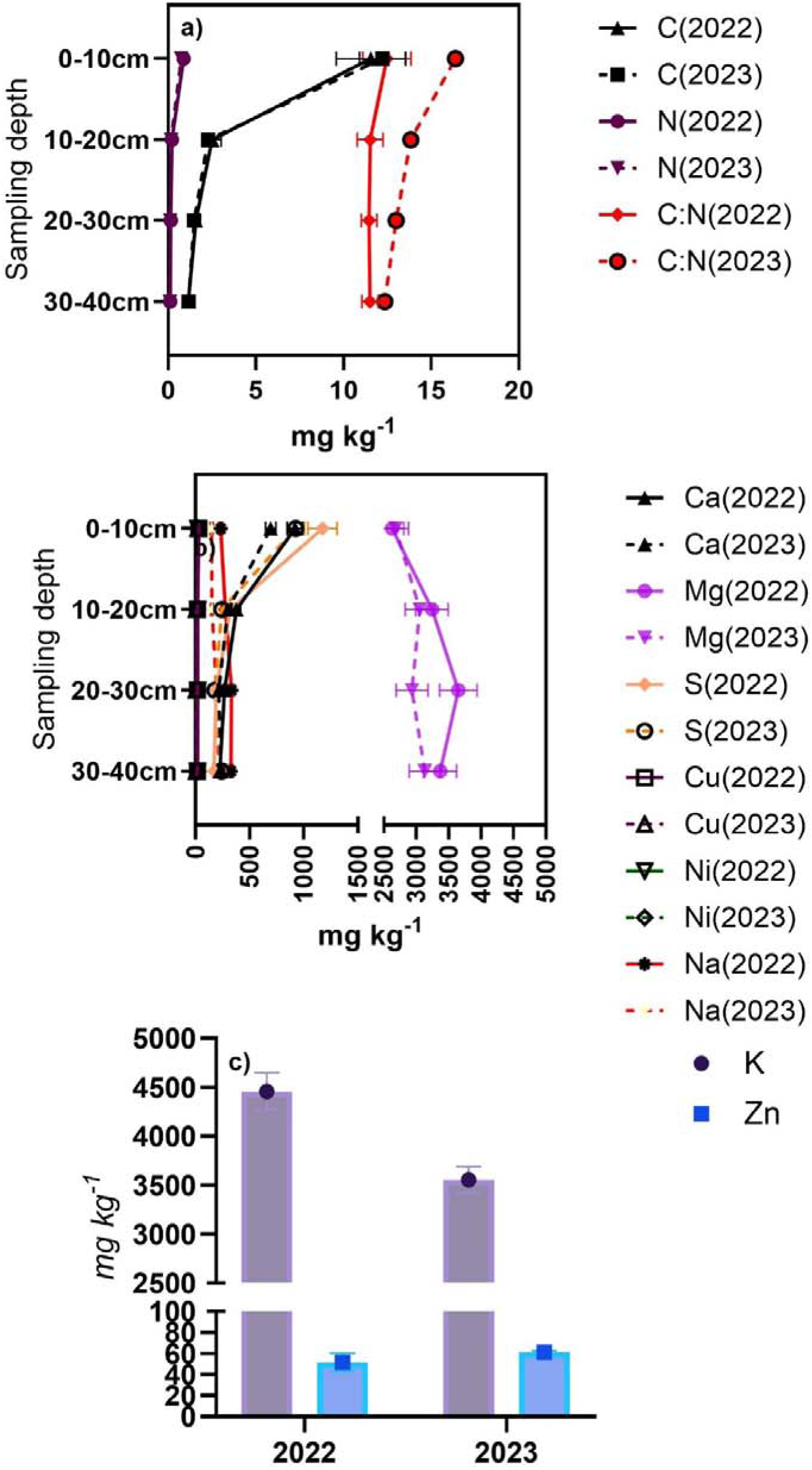
Effect of treatment and measurement period upon a) Total Carbon, Nitrogen and Carbon:Nitrogen ratio, b) Total macro- and micro-nutrients (Ca, Mg, S, Cu, Ni and Na) and c) K and Zn. Only significant effects from linear mixed model (REML) are presented in the figure. Whiskers denote standard errors of means. Data points for which the error bars are not visible, the error bars are smaller than the symbols used for data points.

### 3.4 Seasonal changes in soil microbial community structure

Microbial phenotypic structure determined by phospholipid fatty acid analyses showed significant temporal variations, however the distinct microbial groups (Actinobacteria, Gram-positive bacteria, Gram-negative bacteria, Other bacteria, Fungi, MicroEukaryotes and Unknown fatty acids) did not show significant treatment effect (Figure 7a; Table 4). The microbial community structure was heavily dominated by bacteria over fungi, with clear Gram-positive dominance in the Autumn season, and generic bacterial biomarkers forming the biggest group in all the other seasons. This temporal variations were highly visible in fungi: bacteria (F:B) and Gram positive: Gram negative (G+:G-) ratios (Figure 7b, 7c). Considering actinobacteria also are part of the Gram positive group, they formed the most dominant bacterial group over Gram negative bacteria across treatments and seasons (Figure 7a, 7c). F:B ratio and G+:G- ratio exhibited a mutually inverse temporal trend; F:B ratio was lowest in Autumn and increased over the course of the year to Summer, while G+:G- ratio was highest in Autumn and decreased over the course of the year to Summer. Other fatty acid ratios, such as Cyc17:0 to 16:1n7 (Figure 7d), and saturated fatty acids to monounsaturated fatty acids (Figure 7e) did not show any treatment or season effect (Table 4).

**Table 4:**
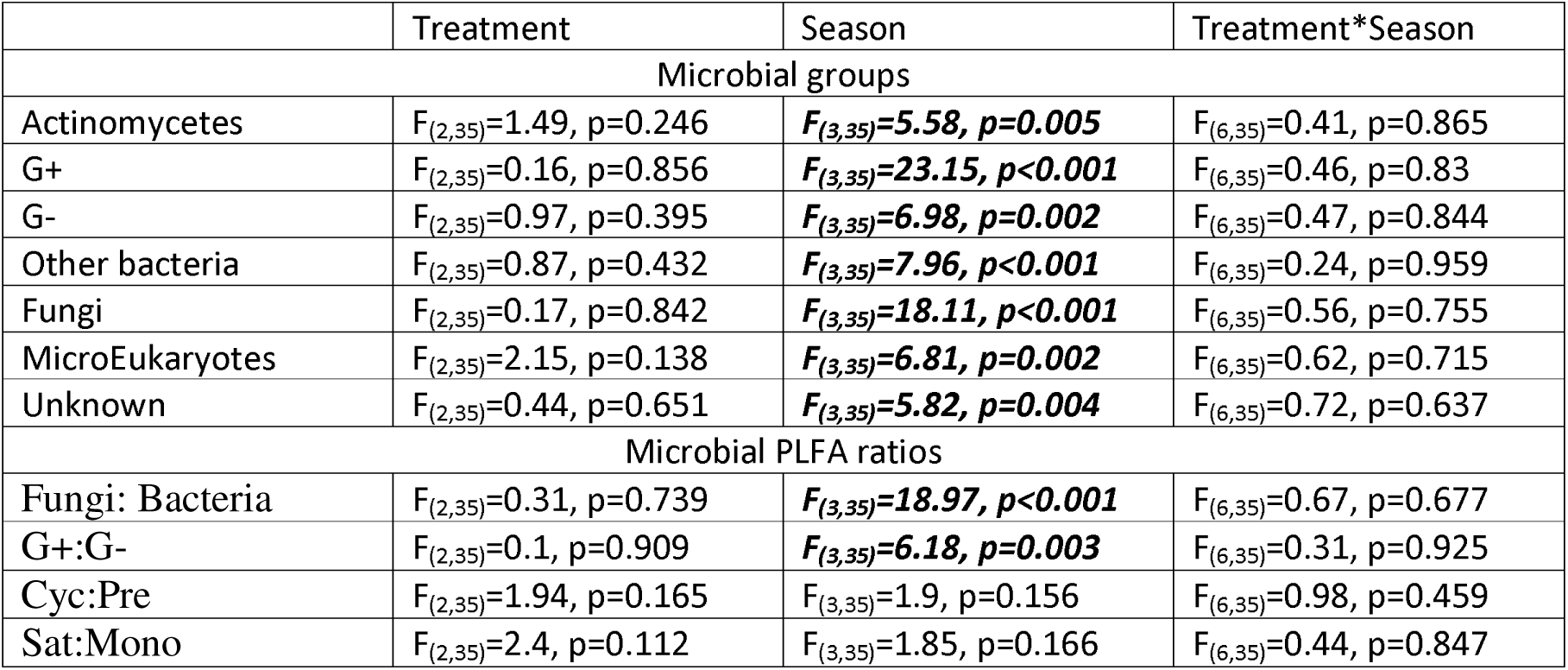
Two way ANOVA for aggregated PLFA data with respect to microbial groups, and ecologically important fatty acid ratios, showing statistical significance of the effects of treatment, season and interactions between treatment and season.

|  | Treatment | Season | Treatment*Season |
| --- | --- | --- | --- |
| Microbial groups |  |  |  |
| Actinomycetes | $F_{(2,35)}=1.49, p=0.246$ | **$F_{(3,35)}=5.58, p=0.005$** | $F_{(6,35)}=0.41, p=0.865$ |
| G+ | $F_{(2,35)}=0.16, p=0.856$ | **$F_{(3,35)}=23.15, p<0.001$** | $F_{(6,35)}=0.46, p=0.83$ |
| G- | $F_{(2,35)}=0.97, p=0.395$ | **$F_{(3,35)}=6.98, p=0.002$** | $F_{(6,35)}=0.47, p=0.844$ |
| Other bacteria | $F_{(2,35)}=0.87, p=0.432$ | **$F_{(3,35)}=7.96, p<0.001$** | $F_{(6,35)}=0.24, p=0.959$ |
| Fungi | $F_{(2,35)}=0.17, p=0.842$ | **$F_{(3,35)}=18.11, p<0.001$** | $F_{(6,35)}=0.56, p=0.755$ |
| MicroEukaryotes | $F_{(2,35)}=2.15, p=0.138$ | **$F_{(3,35)}=6.81, p=0.002$** | $F_{(6,35)}=0.62, p=0.715$ |
| Unknown | $F_{(2,35)}=0.44, p=0.651$ | **$F_{(3,35)}=5.82, p=0.004$** | $F_{(6,35)}=0.72, p=0.637$ |
| Microbial PLFA ratios |  |  |  |
| Fungi: Bacteria | $F_{(2,35)}=0.31, p=0.739$ | **$F_{(3,35)}=18.97, p<0.001$** | $F_{(6,35)}=0.67, p=0.677$ |
| G+:G- | $F_{(2,35)}=0.1, p=0.909$ | **$F_{(3,35)}=6.18, p=0.003$** | $F_{(6,35)}=0.31, p=0.925$ |
| Cyc:Pre | $F_{(2,35)}=1.94, p=0.165$ | $F_{(3,35)}=1.9, p=0.156$ | $F_{(6,35)}=0.98, p=0.459$ |
| Sat:Mono | $F_{(2,35)}=2.4, p=0.112$ | $F_{(3,35)}=1.85, p=0.166$ | $F_{(6,35)}=0.44, p=0.847$ |

**Figure 7:**
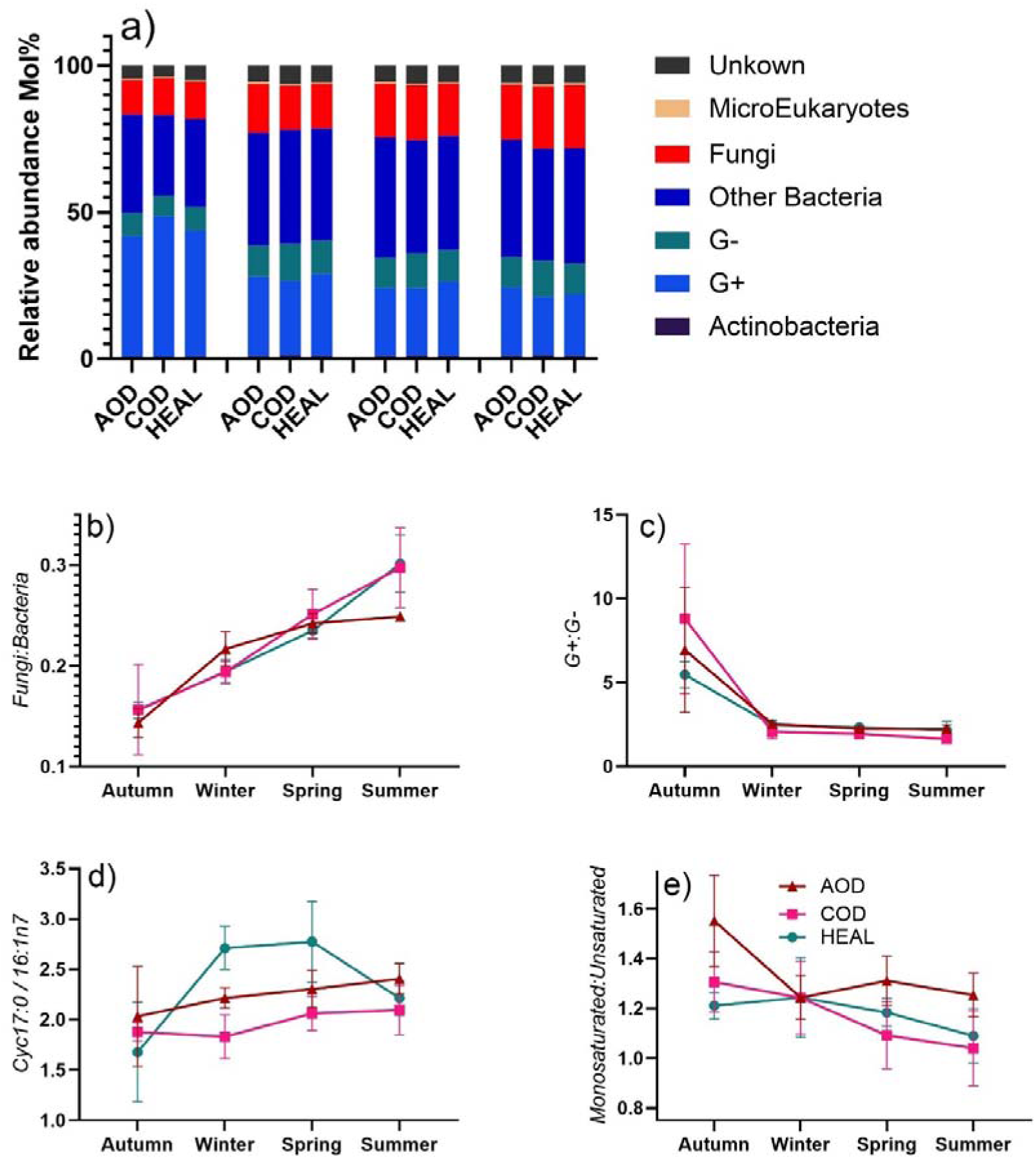
Microbial community structure and fatty acid ratios. a) mean values of relative abundance of different microbial groups as determined by PLFA analyses. Effect of treatment and season (two-way ANOVA) upon b) Fungi:Bacteria ratio, c) G+:G- ratio, d) cyc17:0/16:1n7 ratio and d) Saturated fatty acids:Monounsaturated fatty acids ratio. Whiskers denote standard errors of means. Data points for which the error bars are not visible, the error bars are smaller than the symbols used for data points.

Principal components (PC) analysis showed significant discrimination of both treatment and season. PC 1 and 2 collectively accounted for 71% of the variation (Figure 8). PC1 which accounted for 55.3% of the variation separated Autumn from the rest of the seasons. This separation was characterised by high relative abundance of one G+ fatty acid, namely i15:0 in Autumn (Figure 8c). PC2 which accounted for 15.7% of the variations, separated AOD from the other treatments, and this separation was characterised by a mix of different bacterial fatty acids (Figure 8c). It should be noted that PC1 that accounted for more than half of the variation did not separate the treatments, indicating high similarity in microbial community structure between the treatments.

**Figure 8:**
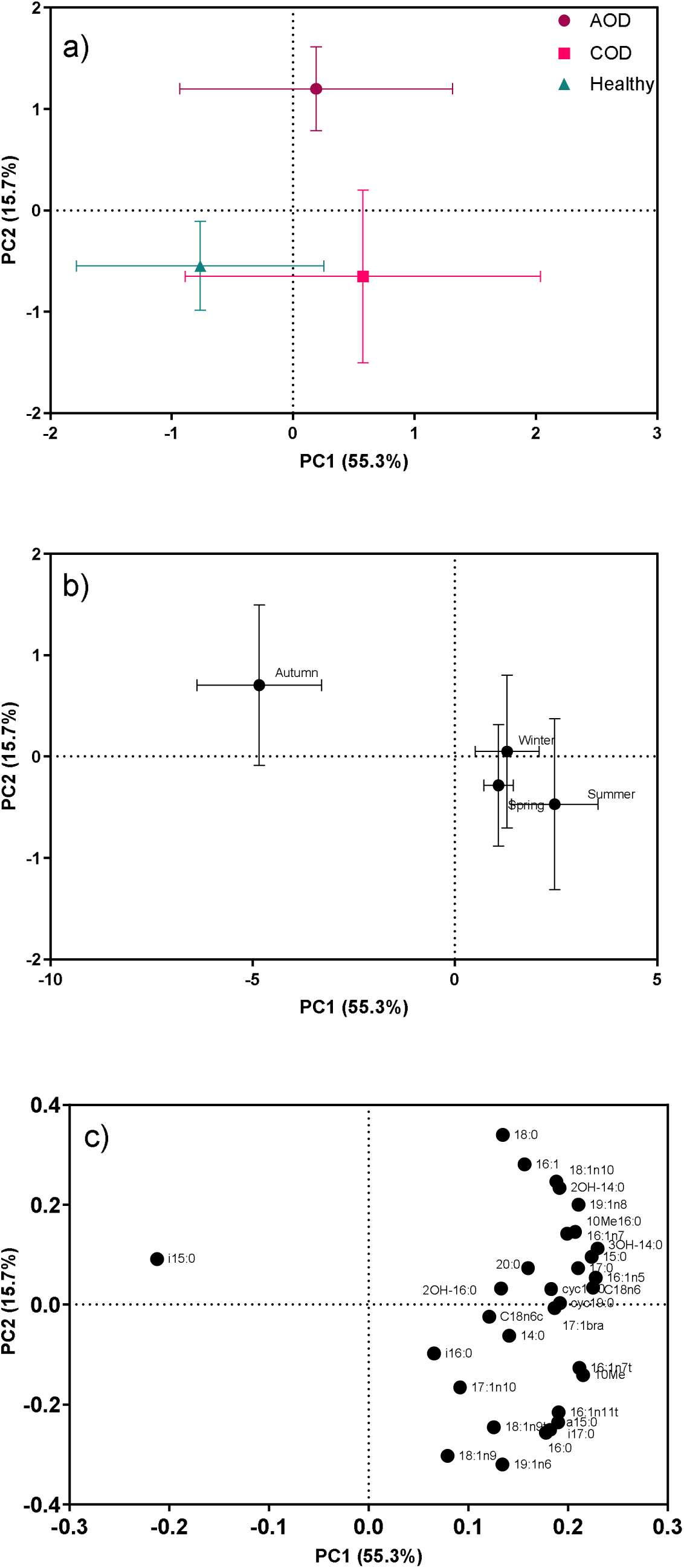
Effects of treatment and season upon phenotypic community structure of soil microbes determined by PLFA analyses, as shown by principal components (PC) analysis. a) ordination of PC1 and PC2 discriminating tree health treatments, b) ordination of PC1 and PC2 discriminating seasons, and c) associated loadings for individual PLFAs. Whiskers denote standard errors of means.

### 3.5 Monthly dynamics of leaf litter quantity and quality

Leaf litter quantity showed significant temporal variation, however there was no significant treatment effect on any of the components of leaf litter (leaves, wood, seeds and flowers). Flowers were found in the litter trap only in the summer period from May to July, and seeds were found in the traps only in late summer (August) and autumn (October) period. The leaf quantity was significantly higher in late autumn and the early winter period, reaching more than 30 g per trap dry weight during the December period. Similarly, Flower dry weight was highest in early summer period, reaching nearly 10 g dry weight per trap at the end of May. Seed dry weight reached the highest of over 5 g per trap in October (Figure 9).

**Figure 9:**
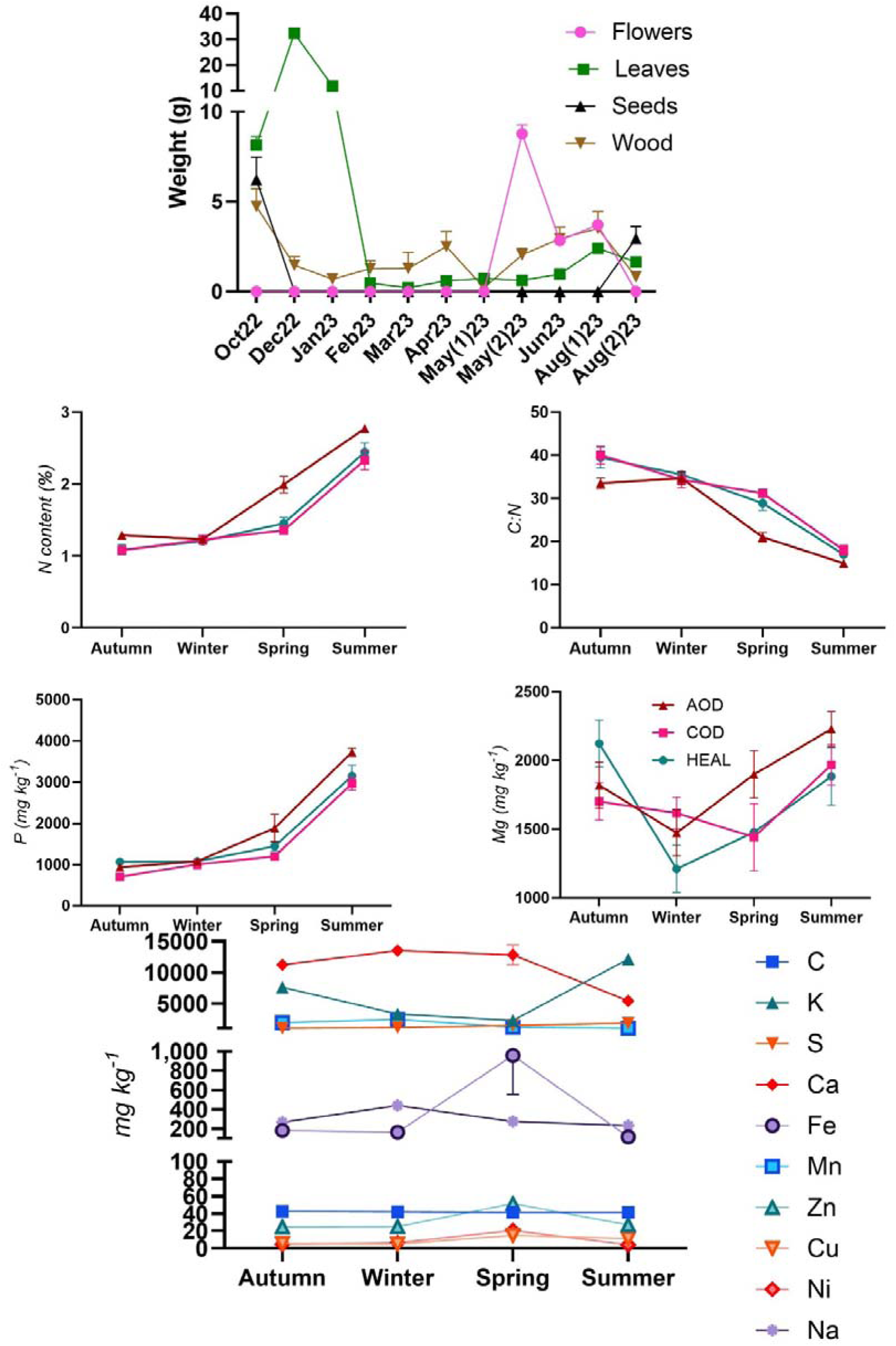
Effect of treatment and measurement period upon a) litter weight (grams per trap) separated as leaves, wood, seed and flowers. Only significant effects from linear mixed model (REML) are presented. Effect of treatment and season upon leaf litter chemistry b) N, c) C:N ratio, d) P, e) Mg, and f) other macro- and micro-nutrients. Only significant effects from two-way ANOVA is presented in figures. Whiskers denote standard errors of means. Data points for which the error bars are not visible, the error bars are smaller than the symbols used for data points.

**Figure 10:**
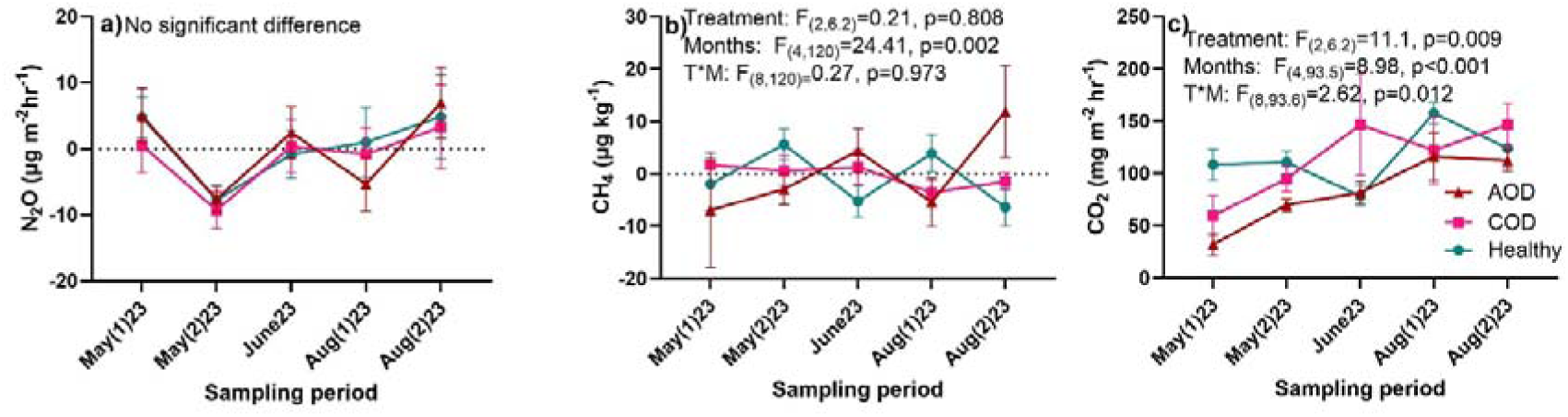
Effect of treatment and measurement period (linear mixed model using REML) upon soil fluxes of a) N_2_O b) CH_4_ and c) CO_2_. Whiskers denote standard errors of means. Data points for which the error bars are not visible, the error bars are smaller than the symbols used for data points.

The pH of tree litter did not differ between treatments and it stayed at mean pH of 4.7 (Table 5). There was no difference in chemistry (nutrient and trace element contents) between treatments for flower and seed samples. However, leaf samples showed significantly greater macronutrient content (N, P, S) and reduced C:N ratio in AOD affected trees especially in spring and summer seasons (Fig 9b, c, & d). Carbon and all the measured macro and micronutrient contents, and pH of the leaf litter also displayed significant seasonal changes (Fig 9e). Similarly for wood litter, pH and all nutrients except S, Mn, Zn showed significant seasonal variations (Supplementary Fig.2; Supplementary Table 3), with generally higher content in wood in the summer season compared to the other seasons. Nutrients such as K and Zn were higher under AOD and COD trees compared to healthy trees showing significant treatment effect (Supplementary Fig.2; Supplementary Table 3).

**Table 5:**
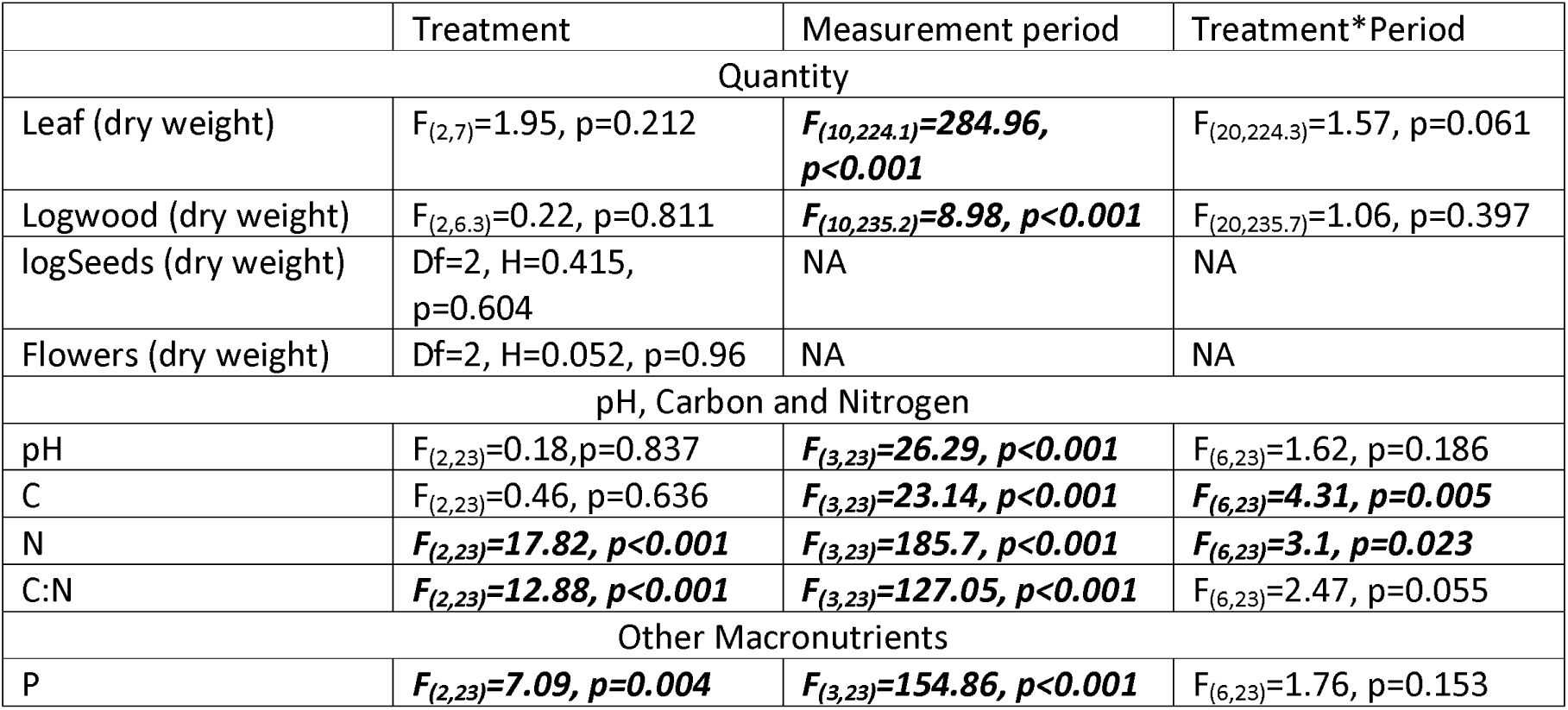
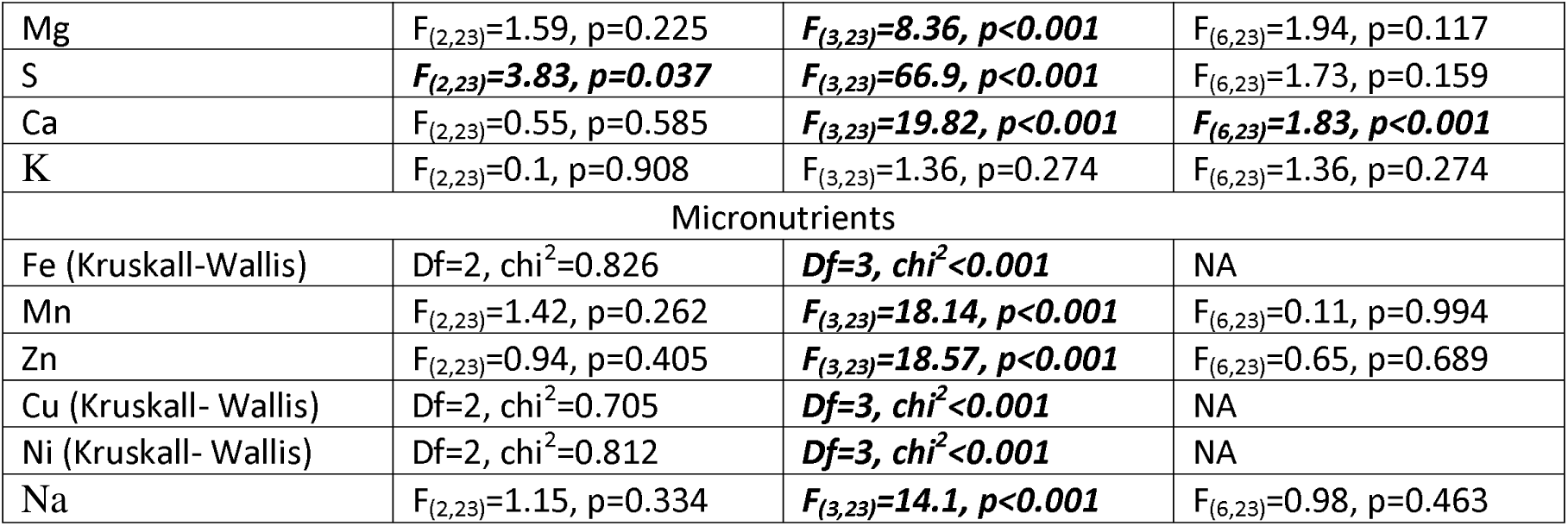
Linear mixed model (REML) for litter quantity and two-way ANOVA for litter chemistry (for leaf component only), showing statistical significance of the effects of treatment (tree health status), depth, and the interactions between treatment and season. Statistically significant figures are presented in bold italics.

### 3.6 Greenhouse Gas (GHG) emissions

N_2_O (mean fluxes ranged from -9.3 to 7 µg m^2^ hr^-1^) and CH_4_ (mean fluxes ranged from -6.9 to 11.9 µg m^2^ hr^-1^) emissions were minimal from oak root zones of any treatment and did not show any treatment effect. Notably highest mean values for both N_2_O and CH_4_ were recorded in AOD root zone soil for August month. CO_2_ emissions made the largest contribution to GHG emissions from oak root zone soil with mean range from 32.2 to 175.5 mg m^2^ hr^-1^. CO emissions also showed significant treatment effect with significantly lower emissions from AOD root zone soil. Both carbon emissions (CO₂ and CH₄) exhibited a significant seasonal effect, characterised by an increase in CO₂ emissions from May (spring) to August (summer) and elevated CH₄ emissions during the August period.

## 4. Discussions

### 4.1 Soil chemistry: Increased nutrient availability under AOD trees, with significant temporal and depth level changes

Soil nutrient availability showed a very strong temporal and soil depth effect. There was also significantly greater macronutrient (Nitrate-N, ammonium-N, P and K) availability in AOD root zone soil, possibly because of the reduced uptake of nutrients by the AOD-affected trees. Some of the micronutrients such as Ni and Na, and trace elements such as Al also showed greater concentration under AOD root zone. It should be noted that the level of total C and total nutrients did not vary between treatments. The difference in nutrient availability is therefore very likely directly influenced by the tree health condition, considering that there was no pre-existing difference in total nutrient or total carbon contents in the root zone soil (Table 3) of these trees at the start of the monitoring period. The findings of elevated nutrient availability for surface soils in this study contrast the findings of Shaw et al. (2025) , who reported lower available P and Mg in AOD-associated soils at 5-15 cm depth. However, their study was a single timepoint assessment taken in July and used different extraction methods (Olsen for P and BaCl_2_ for Mg), which may explain the discrepancy.

Carbon and almost all the measured nutrients in soil showed significant reduction over the monitoring period, indicating potential further loss of nutrients in this nutrient poor woodland. However, it should be noted that the measurement period started in Autumn and ended in Summer, and thus this observation may just be an annual cycle where nutrients get replenished with litter fall in Autumn.

The observed variation in nutrient availability with soil depth and season aligns with expected patterns driven by biological and environmental factors as has been frequently recorded in different forest ecosystems in different soil types (Dhandapani et al., 2023a; Jobbágy and Jackson, 2001; Rahman et al., 2022). Macronutrient inputs which are heavily dependent on leaf litter addition such as N, P, K and Ca are notably greater in the surface layers and decreased with depth. The results emphasise that most of the nutrient activity is in the top 10 cm of the soil, where more frequent four-weekly sampling was conducted in this study.

Autumn had increased nutrient availability, which was also reflected in in-field continuous measurement of soil pore water nitrate concentrations using state of the art real time nitrate sensing (Lu et al., 2024). These continuous measurements showed free nitrate in porewater of an AOD tree’s soil (Location 1 in Lu et al., 2024) was consistently higher than for a healthy tree (Location 2 in Lu et al., 2024). While nitrate levels on both locations dropped to near zero in winter, the nitrate levels remained elevated for 20 days longer for the AOD tree. These continuous measurements over six months align closely with the monthly snapshot sampling and conventional laboratory analyses, confirming the observed seasonal trends. Increased nutrient availability in forest soils during autumn has been previously observed in several different regions (Díaz-Raviña et al., 1995; Farley and Fitter, 1999; Wang et al., 2022) and is likely driven by multiple interacting factors, including plant physiological and soil microbial activity responses to weather conditions. Mild temperatures (Ca. 12° c: Fig. 2) and moderate soil moisture (37.2 %: Fig.2) in autumn create favourable conditions for nutrient release via mineralization of litter and soil organic matter. The rate of fresh leaf litter input starts to increase in early autumn due to the annual leaf drop of deciduous trees, while decomposition of litter deposited in spring and summer, which experiences slower decomposition due to preceding drier conditions, is reactivated by autumn rewetting. Additionally, uptake of nutrients by trees decreases as they enter dormancy in preparation for the winter period, contributing to higher soil nutrient concentrations (Farley and Fitter, 1999). During the period of leaf senescence, trees also re-allocate nutrients from leaves to perennial tissues (Lawrence and Melgar, 2018), reducing reliance on soil nutrient uptake. It is likely that these combined processes explain the elevated soil available nutrient levels in autumn.

### 4.2: Soil microbial communities: Bacterial dominance with significant temporal and AOD effect

The soil microbial communities irrespective of the tree health were heavily dominated by bacteria over fungi. However, generally fungi are more efficient in breaking down complex organic matters in forest ecosystems (Bardgett, 2005; Fierer et al., 2009), and bacteria often use the byproducts from fungal decomposition in these systems (Moore-Kucera and Dick, 2008). It may be the case that there was more fungal activity in woody leaf litter, and the fungal decomposition products left in the soil (sampled in this study) are used by the bacterial communities, aiding heavy dominance of bacteria over fungi in surface soils. This kind of unexpected bacterial dominance in forest soil systems with complex organic matter input is seen in different temperate (Bååth and Anderson, 2003; San Román et al., 2024) and tropical (Dhandapani et al., 2020; Dhandapani et al., 2019b) regions.

The microbial communities showed significant changes with season, however individual phenotypic groups or the fatty acid ratios of ecological significance did not significantly differ between root zone soil of diseased trees and healthy trees. Nevertheless, principal component analyses taking into account the changes in individual fatty acids showed that AOD root zone soil had significantly different microbial communities. One sole fatty acid belonging to the Gram-positive group separated Autumn from the rest of the seasons. This also coincides with increased nutrient availability in Autumn staying distinct from other seasons. It has been observed in several ecosystems that the soil nutrient availability and soil microbiological communities significantly interact with each other (Cao et al., 2024; Dhandapani et al., 2019b).

Within bacterial groups, Gram-positive bacteria generally degrade more complex organic carbon substrates, while Gram-negative bacteria prefer more labile substrates (Fanin et al., 2019). Among Gram-positive bacteria, actinobacteria - a filamentous subgroup - have adaptation to break down complex, woody organic matter, functioning similarly to fungi in this respect (Barka et al., 2016; Bhatti et al., 2017). Hence, in an oak forest systems, where complex woody material constitutes a significant portion of aboveground litter input, Gram-positive bacteria tend to be more abundant than Gram-negative bacteria. The increased abundance of Gram-positive bacteria in Autumn may also relate to greater quantity of carbon and nutrient input through seasonal litter deposition. The abundance of Gram positive bacteria has previously been found to be favoured by higher nitrogen and oxygen availability (Dhandapani et al., 2020; Dhandapani et al., 2019b; Liu et al., 2015; Smith et al., 2014). During Autumn, there was greater nitrogen addition from leaf litter - particularly from AOD trees - and resultant greater nitrogen availability in aerobic surface soils which likely influenced the abundance of Gram-positive bacteria in Autumn compared to other seasons. This interaction is further supported by the opposing seasonal trends in the G+:G- ratio and F:B ratio; the F:B ratio was at its lowest in Autumn, while G+:G- was at its highest suggesting that the decline in fungal abundance during Autumn was driven by the significant increase in G+ relative abundance. This shift might have been influenced by greater available nitrogen concentrations in soil.

Under stressful conditions, microbes modify their cell membrane composition by converting relatively unstable fatty acids such as monoenoic fatty acids (eg. 16:1n7, 18:1n7 etc) to more stable cyclopropane fatty acids (eg. cyc17:0, cyc19:0 etc) to maintain membrane integrity and function (Kaur et al., 2005). It follows that the ratio between cyclopropane fatty acids and their monoenoic precursors is a good indicator for stress experienced by microbial communities in soil systems (Kaur et al., 2005; Wilkinson et al., 2002). It is interesting to note that there was not even a seasonal change in this ratio in our study, despite the strong seasonal changes in relative abundance of different microbial phenotypic groups. This indicates that there are no detectable differences in microbial physiological stress between root zone soil of trees with varying health status or across different seasons (Bossio and Scow, 1998; Kieft et al., 1994; Liu et al., 2015; Wilkinson et al., 2002). Similarly, the ratio of saturated to monounsaturated fatty acids, which serves as an indicator of nutritional stress in soils (Moore-Kucera and Dick, 2008; Zelles et al., 1992) showed no significant variation with season or treatment. This further highlights the absence of additional stress to soil microbial communities associated with acute or chronic oak decline.

### 4.3: Leaf litter chemistry: Significantly higher macronutrient content in litter from AOD trees

Forest litterfall forms an important part of forest biogeochemical cycling, serving as a primary source of essential nutrients that sustain forest ecosystems (Ge et al., 2013). This role is particularly important for deciduous forests, which produce the greatest quantity of litterfall among forest types in temperate regions (Landsberg and Gower, 1997). In our study, there was no difference in litter quantity between the treatments despite the difference in canopy sizes. This may be due to the fact that the litter traps were placed within 1.5m distance from the stem and all the trees generally had canopy cover of the focal tree over the traps. Although litter fall collected in traps is generally considered a measure of litterfall for a given forest area rather than from any specific tree (Ukonmaanaho et al., 2020), the litter in each trap in our study was likely predominantly sourced from the targeted tree. This inference is supported by the sheltered conditions within the woodland interior and recorded absence of wind during each sampling visit, which would limit the movement of litter between trees.

Litter from AOD trees had significantly greater macronutrient content, especially in the summer and spring months. As discussed (section 4.1), this was followed in autumn by elevated nutrient availability in soils beneath AOD-affected trees, possibly due to this increased nutrient addition through their enriched leaf and wood litter. This is very likely due to poor resorption of nutrients by AOD trees before leaf senescence (Hagen-Thorn et al., 2006; Lawrence and Melgar, 2018).

Particularly P and N content were greater in AOD tree leaf litter, and it is known that healthy trees can re-translocate these two macronutrients more efficiently (up to 80%) than any other nutrients further indicating failure in this function in diseased trees (Hagen-Thorn et al., 2006). Nevertheless, macronutrient content in leaf litter is generally high in the summer and spring period for all trees, which also resulted in greatly reduced C:N ratio. This temporal change reflects the seasonal patterns of a deciduous forest. Spring and summer periods are the most active period where nutrients from soil are taken up for tree growth, thus any litterfall during this period had greater macronutrient concentrations than other seasons.

### 4.5: Soil GHG emissions: Low CO_2_ emissions from AOD rhizosphere

N₂O emissions from soils are primarily produced through microbial processes, including nitrification and denitrification (Yu et al., 2022). Both processes are interconnected; the mineralization of organic nitrogen supplies ammonia for nitrification, which then generates nitrate that feeds denitrification under low oxygen conditions. In this study, soil N_2_O emissions were near zero and did not show any changes with season or tree health status. This coincided with very low nitrate-N (monthly mean 66 to 282 mg N kg^-1^ ) and ammonium-N (mean 72 to 210 mg N kg^-1^ ) activity in soil generally for all treatments in spring and summer period, probably due to depletion of soil mineral N pools due to root uptake during active growth. The exception was the Autumn period, where detection of elevated nitrate (1054 mg N kg^-1^) indicated that nitrification may have been active in this season, likely associated with increased nitrogen inputs from litter decomposition and favourable pH and moisture in the litter microenvironment. However, greenhouse gas measurements were made only in Spring and Summer in this study due to logistical issues, limiting our understanding of the direct relationship between N_2_O emissions and seasonal soil nitrate and ammonium activity. Similarly, CH_4_ emissions were also near zero, which is not surprising as CH_4_ is produced as a result of anaerobic decomposition, and the woodland surface soils were never in anaerobic conditions during any of the sampling period. As methanogenic conditions typically develop under strictly anaerobic conditions in waterlogged soils most common in wetlands, peatlands, and flooded soils, where prolonged saturation limits oxygen diffusion, methanogenesis is unlikely to occur in forest surface soil horizons that are not waterlogged. It is also possible that any methane produced in deeper anaerobic layers was consumed by methanotrophs in aerobic surface layers.

CO_2_ which is the most dominant GHG in this woodland showed significant temporal variations and was also affected by the tree health status. The temporal changes followed the changes in soil temperature with season, with CO_2_ emissions increasing with increased temperatures. This temperature dependence of soil CO_2_ emissions is a well observed and established relationship (Conant et al., 2011; Davidson and Janssens, 2006). Total soil CO_2_ emissions consist of two distinct components, autotrophic respiration and heterotrophic respiration (Hergoualc’h and Verchot, 2011; Ryan and Law, 2005). Autotrophic respiration, driven by root activity, is part of the immediate photosynthetic cycle and, as such, does not contribute to soil carbon loss. In contrast, heterotrophic respiration arises from microbial decomposition, which directly contributes to soil carbon loss (Dhandapani et al., 2019a). As the GHG emissions were measured within 1.5 m of mature oak tree stems and therefore well within the root-influenced zone, it is certain that autotrophic respiration significantly contributed to total soil CO_2_ emissions. There was no indication of reduced microbial decomposition in the AOD root zone soil. These soils generally had higher nutrient activity and minimal changes to microbial community structure compared to the root zone soil of healthy trees. Autotrophic emissions, which contribute approximately 50% of total soil emissions in forests, are mainly controlled by tree physiology, and can be greatly impacted by tree health and disease status (Murdiyarso et al., 2017; Ryan and Law, 2005). Thus, it is likely that reduced autotrophic root respiration in AOD-affected trees contributed to the observed reduction of total soil CO_2_ emissions.

Even though both the carbon and nitrogen contents of the soil were reduced over time for the duration of the monitoring period (Fig 6), C:N ratio increased because of the relatively greater loss of nitrogen. The lack of N_2_O emissions indicate that most of this nitrogen loss perhaps occur through leaching. This changes in soil C:N ratio may have significant influence in soil and ecosystem functioning in this oak woodland (Qi et al., 2022).

## 5. Synthesis and Conclusions

Taken together, our study shows that AOD-affected trees exhibit impaired physiological nutrient management compared to COD and healthy trees. This is evidenced by higher macronutrient concentrations in AOD litter, suggesting reduced nutrient resorption and re-allocation before leaf senescence. Additionally, AOD-affected trees had a less active rhizosphere, indicated by reduced soil CO_2_ emissions and increased nutrient availability. In terms of seasonal changes, the Autumn was distinct with greater nutrient availability, very possibly from greater litter nutrient addition from the preceding summer. This coincided with changes in microbial communities with greatly increased Gram-positive relative abundance in Autumn. This seasonal difference was particularly visible under AOD trees, that were shedding litter with greater nutrient content in spring and summer, and also had greater nutrient content in soil than COD and HEAL trees. In addition to greater nutrient input via litter, the greater quantity of available nutrients in soil is likely also due to poor uptake by root systems linked to impaired vascular transport as a result of the activity of AOD pathogens. The lower CO_2_ emissions further suggest decline in autotrophic contributions from the rhizosphere, reinforcing the notion of a less root activity in AOD affected trees.

This study is the first of its kind to comprehensively monitor the temporal biogeochemical dynamics of root zone soil of oak trees of different health statuses. It examined the total content and availability of all the essential macro- and micro-nutrients and trace elements in the soil. Additionally, the study tracked nutrient inputs through different litter components, outputs through GHG emissions and examines microbial community dynamics over a period of a year. It is clear from our results that biogeochemical dynamics in root zone soil of AOD trees are distinct from that of healthy or COD trees.

Although these findings provide important insight into biogeochemical dynamics in oak decline, this field observation does not disentangle tree scale predisposing causes from the consequences of the AOD decline disease, which are debated but not yet fully understood. The monitored trees had been affected by AOD for several years prior to the study, suggesting that the observed dynamics are likely dominated by the effects of acute oak decline rather than its onset. Nevertheless, this study provides detailed insight into the monthly, seasonal and depth level changes in soil biogeochemistry over a period of a year. It thus provides a baseline for further research to disentangle the causes and effects of AOD, a disease increasingly threatening ecologically and culturally important oak trees in Britain and wider Europe.

## Acknowledgements

We thank Writtle Forest Consultancy for granting access to their woodlands. We are thankful to Anne Dudley, Marta O’Brien, Conor Cooper, Manesha Bongso, Ellie Barbrook, Zuzanna Karolewska, Rory Williams-Burrel, Dr Harry Frost and Dr Peter Jackson for their help with lab analyses. We are also thankful to Rosy Scholes, Sophie Cunnington, TingTing, Camila Beatriz Garcia, Joris Rockx, Monisha Victor Gnanamuthu, Gavkhar Mahmudova for on-field assistance. The corresponding author is also grateful to all the colleagues in The School of Archaeology, Geography and Environmental Sciences in the University of Reading for supporting the author through personally and professionally hard time.

## Author Contributions

SD: Conceptualisation, Investigation, Methodology, Project Administration, Data Curation, Formal Analyses, Visualisation, Writing- Original Draft, Writing- Review and Editing. TM: Formal Analyses, Writing- Review and Editing. BL: Writing- Review and Editing. OB: Writing- Review and Editing. LC: Methodology, Writing- Review and Editing. JL: Writing- Review and Editing, AN: Funding Acquisition, Writing- Review and Editing. XN: Funding Acquisition, Writing- Review and Editing. TR: Funding Acqisition, Writing- Review and Editing. JJ: Writing- Formal Analyses, Review and Editing. HL: Formal Analyses, Writing- Review and Editing. AAA: Formal Analyses, Writing- Review and Editing. LS: Conceptualisation, Funding Acquisition, Methodology, Investigation, Project Administration, Writing- Review and Editing.

