## Supplementary Information for "Forest biogeochemical monitoring indicates altered microbial communities, macronutrient availability, CO_2_ emissions and litter chemistry in root zone soil of oak trees with Acute Decline symptoms"

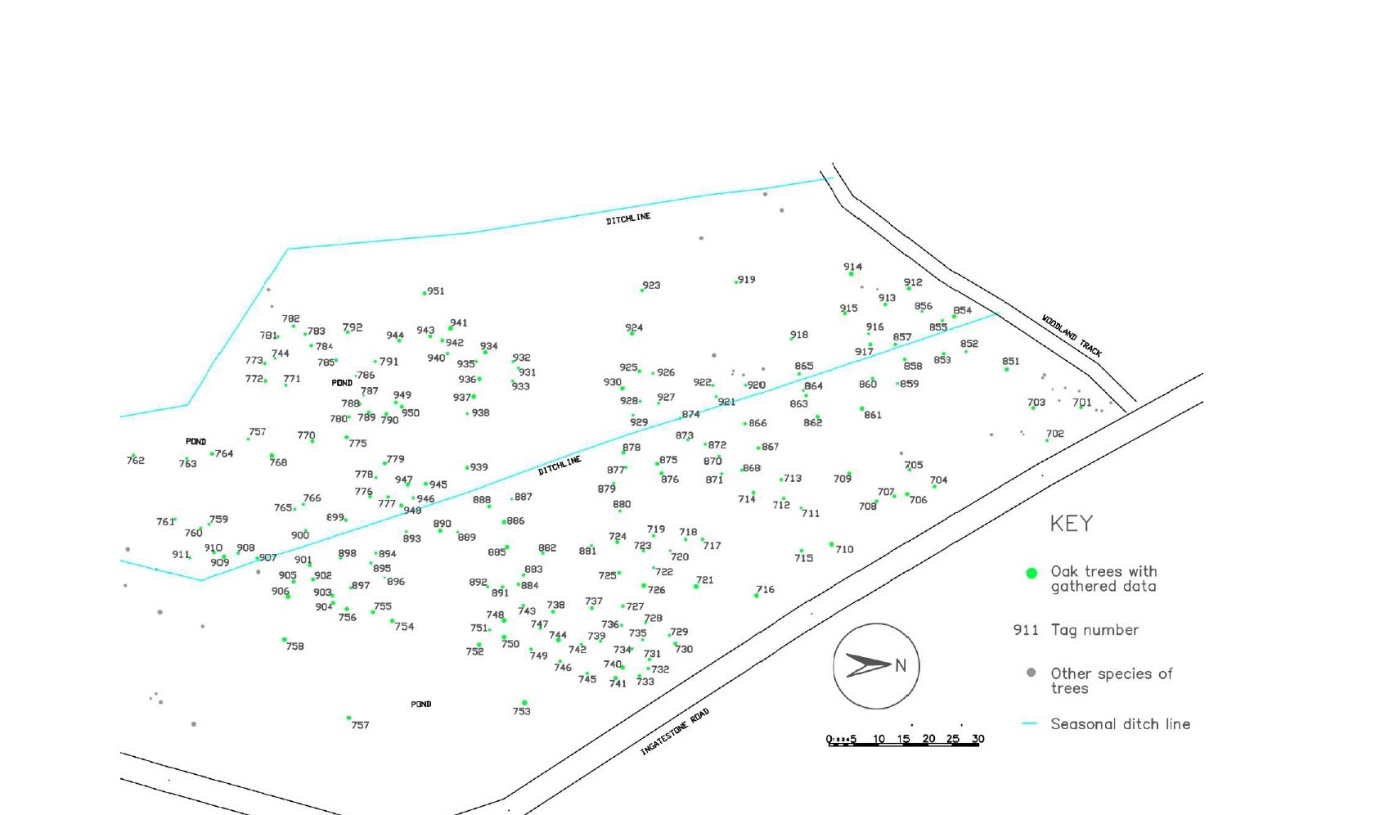

Supplementary Figure 1: Tree map of Writtle woods.

Supplementaty Table 1: Details of the observed historic symptoms in experimental trees used for this study.

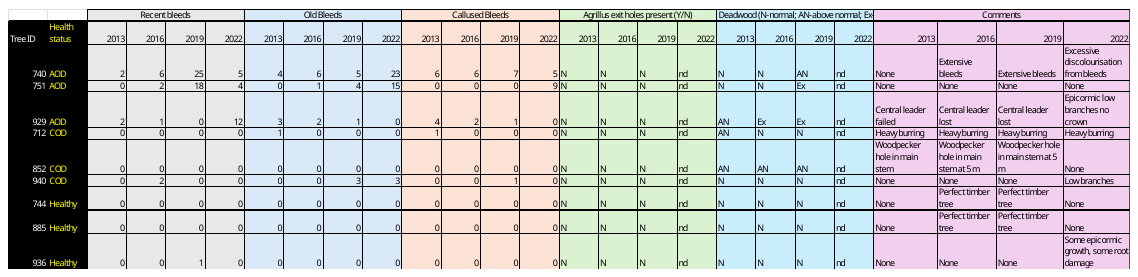

Supplementary Table 2: Sampling dates

| Sampling | Date | Classification |
| --- | --- | --- |
| 1 | 18^th^ and 19^th^ October 2022 | Autumn Quarterly Sampling (0-40 cm soil) |
| 2 | 12^th^ November 2022 | Monthly Sampling (0-10 cm soil) |
| 3 | 13^th^ December 2022 | Monthly Sampling (0-10 cm soil) |
| 4 | 10^th^ January 2023 | Winter Quarterly Sampling (0-40 cm soil) |
| 5 | 14^th^ February 2023 | Monthly Sampling (0-10 cm soil) |
| 6 | 7^th^ March 2023 | Monthly Sampling (0-10 cm soil) |
| 7 | 3^rd^ and 4^th^ April 2023 | Spring Quarterly Sampling (0-40 cm soil) |
| 8 | 3^rd^ May | Monthly Sampling (0-10 cm soil) |
| 9 | 31^st^ May | Monthly Sampling (0-10 cm soil) |
| 10 | 27^th^ June | Monthly Sampling (0-10 cm soil) |
| 11 | 2^nd^ and 3^rd^ August | Summer Quarterly Sampling (0-40 cm soil) |
| 12 | 22^nd^ August | Monthly Sampling (0-10 cm soil) |

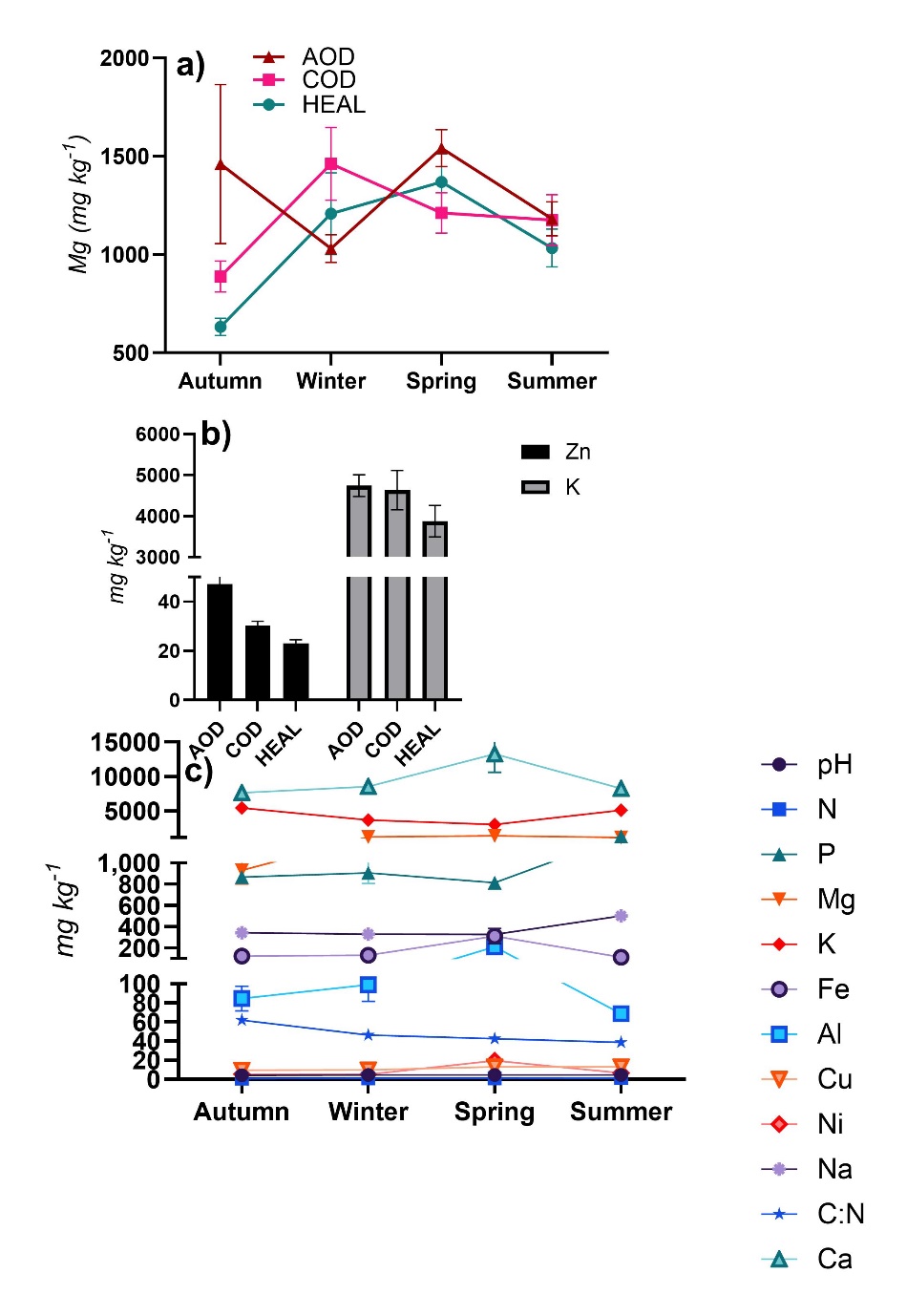

Supplementary Figure 2: Effect of treatment and measurement period upon a)Mg, b) Zn and K, c) C:N ratio, d) P, e) Mg, and c) pH and other macro- and micro-nutrients. Only significant effects from two-way ANOVA are presented in figures. Whiskers denote standard errors of means. Data points for which the error bars are not visible, the error bars are smaller than the symbols used for data points.

Supplementary Table 3: Two-way ANOVA for litter chemistry (for wood component only), showing statistical significance of the effects of treatment (tree health status), depth, and the interactions between treatment and season. Statistically significant figures are presented in bold italics.

|  | Treatment | Measurement period | Treatment*Period |
| --- | --- | --- | --- |
| pH | F_(2,20)_=3.37,p=0.057 | ***F_(3,20)_=6.94, p=0.003*** | F_(6,20)_=0.66, p=0.684 |
| C | F_(2,19)_=0.76, p=0.484 | F_(3,19)_=0.91, p=0455 | F_(6,19)_=0.46, p=0.831 |
| N | F_(2,19)_=0.98, p=0.394 | ***F_(3,19)_=20.86, p<0.001*** | F_(6,19)_=1.85, p=0.142 |
| C:N | F_(2,19)_=1.66, p=0.216 | ***F_(3,19)_=30.75, p<0.001*** | F_(6,19)_=2.12, p=0.098 |
| P | F_(2,20)_=2.89, p=0.079 | ***F_(3,20)_=13.89, p<0.001*** | F_(6,20)_=1.20, p=0.346 |
| Mg | F_(2,20)_=2.60, p=0.099 | ***F_(3,20)_=5.55, p=0.006*** | ***F_(6,20)_=3.36, p=0.019*** |
| S | F_(2,20)_=0.72, p=0.5 | F_(3,20)_=0.84, p=0.489 | F_(6,20)_=0.31, p=0.923 |
| Ca | F_(2,20)_=3.20, p=0.062 | ***F_(3,20)_=3.85, p=0.025*** | F_(6,20)_=1.27, p=0.315 |
| K | ***F_(2,20)_=4.11, p=0.032*** | ***F_(3,20)_=17.99, p<0.001*** | F_(6,20)_=1.4, p=0.262 |
| Fe | F_(2,20)_=0.29, p=0.754 | ***F_(3,20)_=5.09, p=0.009*** | F_(6,20)_=0.15, p=0.988 |
| logMn | F_(2,20)_=1.82, p=0.188 | F_(3,20)_=0.46, p=0.713 | F_(6,20)_=0.31, p=0.926 |
| Zn (Kruskall-Wallis) | ***Df=2, chi^2^=0.049*** | Df=3, chi^2^=0.578 | NA |
| Cu (Kruskall- Wallis) | Df=2, chi^2^=0.945 | ***Df=3, chi^2^=0.021*** | NA |
| Ni (Kruskall- Wallis) | Df=2, chi^2^=0.692 | ***Df=3, chi^2^=0.003*** | NA |
| Na | F_(2,20)_=3.35, p=0.056 | ***F_(3,20)_=7.67, p=0.001*** | F_(6,23)_=1.20, p=0.348 |
